# Virtual Transcriptomics from Routine Histopathology Reveals Metastasis-Associated Programs and Molecular Risk States in Triple-Negative Breast Cancer

**DOI:** 10.64898/2026.11.25.769294

**Authors:** Abay Suleimenov, Darcie D Seachrist, Marlo Nicolas, Ruth Keri, Mihriban Karaayvaz, Murat Yildirim

## Abstract

**Background:** Triple-negative breast cancer (TNBC) is a highly aggressive and biologically heterogeneous disease for which reliable biomarkers of metastatic progression remain limited. Although transcriptomic profiling can reveal clinically important molecular programs, its routine implementation is constrained by cost, tissue availability, and workflow complexity. We hypothesized that metastasis-associated transcriptional states are encoded within routine hematoxylin and eosin (H&E) histopathology and can be recovered using artificial intelligence–enabled virtual transcriptomics.

**Methods:** We applied the Path2Space foundation model to infer genome-wide gene expression from treatment-naïve TNBC H&E tissue microarrays and identified virtual transcriptomic programs associated with lymph node metastasis. We systematically validated candidate genes across three independent international TNBC cohorts (TCGA-BRCA, SCAN-B, and FUSCC). Candidate genes were evaluated in three independent international TNBC cohorts using experimentally measured RNA sequencing, followed by TNBC-specific co-expression network analysis, molecular subtype characterization, and cross-cohort meta-analysis.

**Results:** Virtual transcriptomic analysis nominated 118 candidate genes associated with nodal status at a nominal discovery threshold; these candidates were subsequently evaluated using experimentally measured transcriptomic data across three independent international TNBC cohorts. Cross-cohort validation converged on a parsimonious two-gene Virtual Metastatic Prediction Score (VMPS) composed of TMEM184A and ZNF57, the only genes that remained significantly associated with nodal metastasis across all four datasets. TMEM184A and ZNF57 represented complementary biological programs: a TMEM184A-centered epithelial remodeling network and a ZNF57-centered KRAB zinc-finger transcriptional regulatory network. These transcriptional modules were biologically coherent and demonstrated reproducible associations with established TNBC molecular subtypes. Across TCGA-BRCA, SCAN-B, and FUSCC, the TMEM184A module showed a pooled OR of 1.44 per SD (95% CI, 1.25–1.67; I²=0%), while the two-gene VMPS showed a pooled OR of 1.58 (95% CI, 1.28–1.94). Moreover, higher VMPS and TMEM184A module activity defined reproducible metastatic risk gradients and enriched for lymph node-positive disease across independent cohorts.

**Conclusions:** AI-enabled virtual transcriptomics transforms routine H&E histopathology into a scalable platform for molecular discovery in TNBC. These findings establish virtual transcriptomics as a scalable discovery framework for recovering biologically interpretable, metastasis-associated molecular programs from routine histopathology, with potential applications in precision risk stratification.

## INTRODUCTION

Triple-negative breast cancer (TNBC), defined by the absence of estrogen receptor (ER), progesterone receptor (PR), and human epidermal growth factor receptor 2 (HER2) expression, accounts for approximately 15-20% of breast cancers and represents one of the most clinically aggressive breast cancer subtypes^1–3^. Although historically treated as a single disease, bulk transcriptomic profiling has established that TNBC comprises multiple molecular subtypes with distinct transcriptional programs, biological behavior, and therapeutic vulnerabilities^1,2,4–7^. Despite this molecular heterogeneity and recent advances in systemic therapy, metastatic progression remains the leading cause of TNBC-related mortality^8,9^, creating a need for biomarkers that identify metastasis-prone tumors at the time of diagnosis. Treatment-naive diagnostic specimens provide a unique opportunity to study metastatic programs before they are altered by systemic therapy, which can remodel dissemination programs and prometastatic microenvironments^10–12^. In parallel, federated multi-institutional studies have shown that hematoxylin and eosin (H&E)-based artificial intelligence (AI) can accurately predict histologic response to neoadjuvant chemotherapy in TNBC^13^, highlighting the ability of routine H&E images to capture clinically relevant features of treatment-naïve TNBC.

Recent advances in digital pathology have extended deep learning beyond image classification toward molecular inference. Deep learning models can recover tumor class, actionable genomic alterations, and molecular phenotypes directly from H&E images^14–15^. More recently, a breast cancer-specific study demonstrated that quantitative histologic analysis of epithelial, stromal, immune and spatial tissue features provides robust prognostic information^16^, while pathology foundation models have learned increasingly generalizable morphological representations across tissue types and analytical tasks^17^. Together, these advances demonstrate that routine histopathology contains biologically meaningful information that can be computationally decoded to infer molecular phenotypes directly from tissue morphology.

Spatial and single-cell profiling have clarified why tissue morphology is so informative. Spatial transcriptomics and integrated single-cell/spatial approaches established that tissue architecture is directly linked to location-resolved transcriptional programs^18,19^. In breast cancer, high-dimensional single-cell and imaging-based atlases further demonstrated that tumors are organized not only by cellular composition but also by multicellular neighborhoods and spatial interactions associated with genomic alterations, stromal activation, and clinical outcome^20,21^. In TNBC, multicellular spatial organization has been shown to predict response to immunotherapy^22^, while longitudinal integration of single-cell transcriptomics and spatial profiling has identified distinct response trajectories during checkpoint blockade therapy^23^. More broadly, multimodal spatial atlases of metastatic breast cancer revealed conserved spatial expression programs associated with epithelial plasticity and immune organization across diverse metastatic lesions^24^. Collectively, these studies demonstrate that tissue architecture encodes biologically meaningful spatial programs that shape tumor evolution, therapeutic response, and metastatic progression. Recent mechanistic studies further show that metastatic colonization is governed by distinct spatial architectures, tumor-stromal interactions, and developmental morphogenetic programs^25,26^, providing a strong rationale for recovering these molecular states directly from routine histology.

Emerging virtual spatial transcriptomic approaches now make it possible to recover spatial molecular states directly from routine histology. Emerging multimodal models such as OmiCLIP and STPath, together with systematic benchmarking studies, indicate that spatial gene expression can increasingly be inferred from H&E at scale^27–29^. Building on these advances, Path2Space predicts spatial gene expression directly from breast cancer histopathology slides, enabling large-scale characterization of tumor and microenvironmental states that are otherwise difficult to measure in retrospective clinical cohorts^30^.

In the present study, we leveraged Path2Space to infer genome-wide virtual transcriptomic profiles directly from routine H&E-stained sections of treatment-naïve TNBC tissue microarrays, enabling systematic identification of molecular programs associated with lymph node metastasis. We subsequently evaluated candidate genes across three independent international TNBC cohorts (TCGA-BRCA, SCAN-B, and FUSCC) to identify reproducible metastasis-associated genes across diverse patient populations and sequencing platforms. This cross-cohort framework converged on a two-gene Virtual Metastatic Prediction Score (VMPS) composed of TMEM184A and ZNF57, which consistently associated with lymph node metastasis and demonstrated robust performance across independent cohorts. Systems-level analyses further revealed that these genes represent complementary transcriptional networks linked to epithelial remodeling, cell-surface signaling, and KRAB zinc-finger-mediated transcriptional regulation, providing biological context for metastatic progression. Together, our findings establish a framework in which routine histopathology can be transformed into a source of virtual molecular information, enabling biologically interpretable biomarker discovery and scalable molecular risk stratification without requiring transcriptomic profiling.

## RESULTS

### AI-enabled virtual transcriptomics identifies metastasis-associated genes and places them within TNBC-specific transcriptional programs

To identify molecular programs associated with regional lymph-node involvement in treatment-naïve triple-negative breast cancer (TNBC), we assembled 42 technically available primary tumor tissue microarray (TMA) cases, including 17 lymph node-positive (LN1+) and 25 lymph node-negative (LN0) tumors **(Figure 1A, Supplementary Table S1)**. After image and patient-level curation, 40 cases (16 LN1+ and 24 LN0) were retained for gene-level analyses. Rather than performing RNA sequencing on these specimens, we applied the recently developed Path2Space^30^ framework to infer genome-wide AI-derived virtual transcriptomic profiles directly from routine H&E-stained tissue sections **(Figure 1B)**. Briefly, whole-core images underwent field-of-view matching, tile extraction, stain normalization, feature extraction using the CTransPath pathology foundation model^31^, and multiple-instance learning (MIL)-based prediction of gene expression. Tile-level predictions were aggregated to generate patient-level virtual transcriptomic profiles.

**Figure 1.**
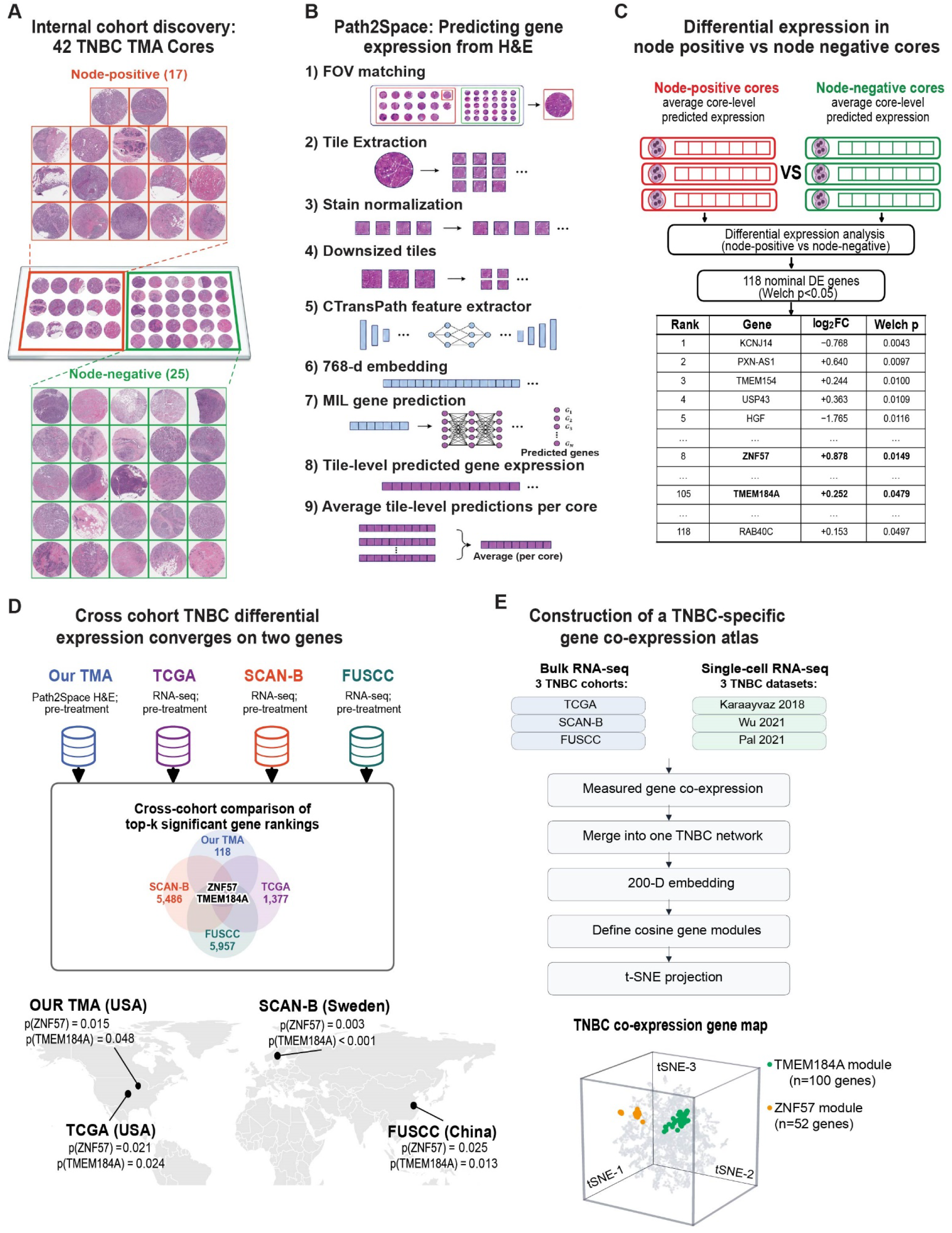
AI-enabled virtual transcriptomics identifies reproducible metastasis-associated genes and maps them to TNBC-specific transcriptional programs. **(A)** Overview of the internal treatment-naïve TNBC discovery cohort comprising 42 tissue microarray (TMA) cores, including 17 lymph node-positive and 25 lymph node-negative primary tumors. **(B)** Path2Space workflow for AI-enabled virtual transcriptomic inference from routine H&E histology. Tissue cores underwent field-of-view (FOV) matching, tile extraction, stain normalization, feature extraction using the CTransPath pathology foundation model, generation of 768-dimensional image embeddings, and multiple-instance learning (MIL)-based prediction of gene expression. Tile-level predictions were aggregated to generate core-level virtual transcriptomic profiles. **(C)** Differential expression analysis comparing predicted virtual transcriptomes between lymph node-positive and lymph node-negative tumors identified 118 nominally significant genes (Welch’s P < 0.05), including the metastasis-associated candidates ZNF57 and TMEM184A. **(D)** Cross-cohort validation across independent treatment-naïve TNBC cohorts from TCGA-BRCA (United States), SCAN-B (Sweden), and FUSCC (China). Cross-cohort comparison of the 118 differentially expressed genes identified TMEM184A and ZNF57 as the only candidates that reproduced consistently across all four independent cohorts spanning distinct patient populations, sequencing platforms, and geographic regions. **(E)** Construction of a TNBC-specific gene co-expression map using measured transcriptomic data from three bulk RNA-sequencing cohorts (TCGA, SCAN-B, and FUSCC) and three epithelial single-cell RNA-sequencing datasets (Karaayvaz et. al^32^, Wu et. al^33^, and Pal et. al^34^). Per-dataset co-expression networks were integrated into a unified confidence-weighted TNBC network, embedded into a 200-dimensional latent space, and used to define cosine-based gene modules. No separate clustering algorithm was applied. The resulting network was visualized using a three-dimensional t-SNE projection, revealing distinct transcriptional neighborhoods centered on TMEM184A (100-gene module) and ZNF57 (52-gene module), thereby providing a systems-level framework for identifying metastasis-associated transcriptional programs beyond individual genes.

Differential expression analysis of the 40 curated predicted transcriptomes identified 118 nominally significant genes associated with regional lymph-node metastasis (Welch’s *P* < 0.05; **Figure 1C, Supplementary Table S2**). These genes were carried forward as candidate metastasis-associated genes for independent validation, providing an unbiased transition from AI-enabled virtual transcriptomic discovery to cross-cohort evaluation. The complete ranked list of all 118 candidate genes is provided in Supplementary Table S2.

We subsequently compared these 118 candidate genes across three independent treatment-naïve TNBC bulk RNA-sequencing cohorts - TCGA-BRCA (United States), SCAN-B (Sweden), and FUSCC (China) – to identify transcriptional signals that generalized beyond the discovery cohort **(Figure 1D)**. Despite substantial differences in cohort composition, sequencing platforms, and geographic origin, TMEM184A and ZNF57 were the only genes consistently associated with lymph node status across all four datasets.

To place these reproducible candidates into their broader biological context, we next constructed a TNBC-specific gene co-expression map using measured transcriptomic data from three bulk RNA-sequencing cohorts (TCGA, SCAN-B, and FUSCC) together with three independent epithelial single-cell RNA-sequencing atlases^32–34^ (**Figure 1E; Supplementary Figure S1**). We derived the network entirely from experimentally measured TNBC co-expression relationships, integrating bulk and single-cell datasets into a unified confidence-weighted graph comprising 513,122 weighted edges connecting 12,609 genes. The resulting graph was embedded into a 200-dimensional latent space, from which transcriptional neighborhoods were identified using cosine similarity (**Supplementary Figures S1 and S2**). Importantly, neither the embedding dimension nor the cosine-similarity threshold represented a locally optimized parameter; both were selected from broad performance plateaus where graph reconstruction, module stability, and biological discrimination remained essentially unchanged (**Supplementary Figure S2**).

Cosine-based neighborhood analysis identified two distinct neighbor sets centered on TMEM184A (100 genes excluding the hub) and ZNF57 (52 genes excluding the hub); the corresponding hub-inclusive module scores contained 101 and 53 genes, respectively. This separation indicates that the two biomarkers occupy distinct regions of the TNBC transcriptional landscape and represent complementary transcriptional programs (**Figure 1E**). Independent biological evaluation further showed that the reconstructed co-expression map recovered established biological gene sets despite being generated without prior pathway annotations or interaction databases such as STRING (**Supplementary Figure S3**). Likewise, the TMEM184A and ZNF57 neighbor sets were consistently identified across bootstrap resampling, embedding dimensions, and similarity thresholds, supporting their stability under the evaluated parameter choices (**Supplementary Figure S2**).

### Cross-cohort validation converges on two reproducible metastasis-associated genes

To determine whether the metastasis-associated genes identified by AI-derived virtual transcriptomics reflected reproducible biology rather than cohort-specific findings, we systematically evaluated their conservation across three independent treatment-naïve TNBC cohorts with measured RNA sequencing and available lymph node status (TCGA-BRCA, SCAN-B, and FUSCC). Differential expression analysis of the internal discovery cohort identified 118 genes associated with nodal metastasis (Welch’s *P* < 0.05), comprising 103 genes with increased and 15 genes with decreased predicted expression in lymph node-positive tumors **(Figure 2A)**. Among the most significantly upregulated genes were ZNF57 and TMEM184A.

**Figure 2.**
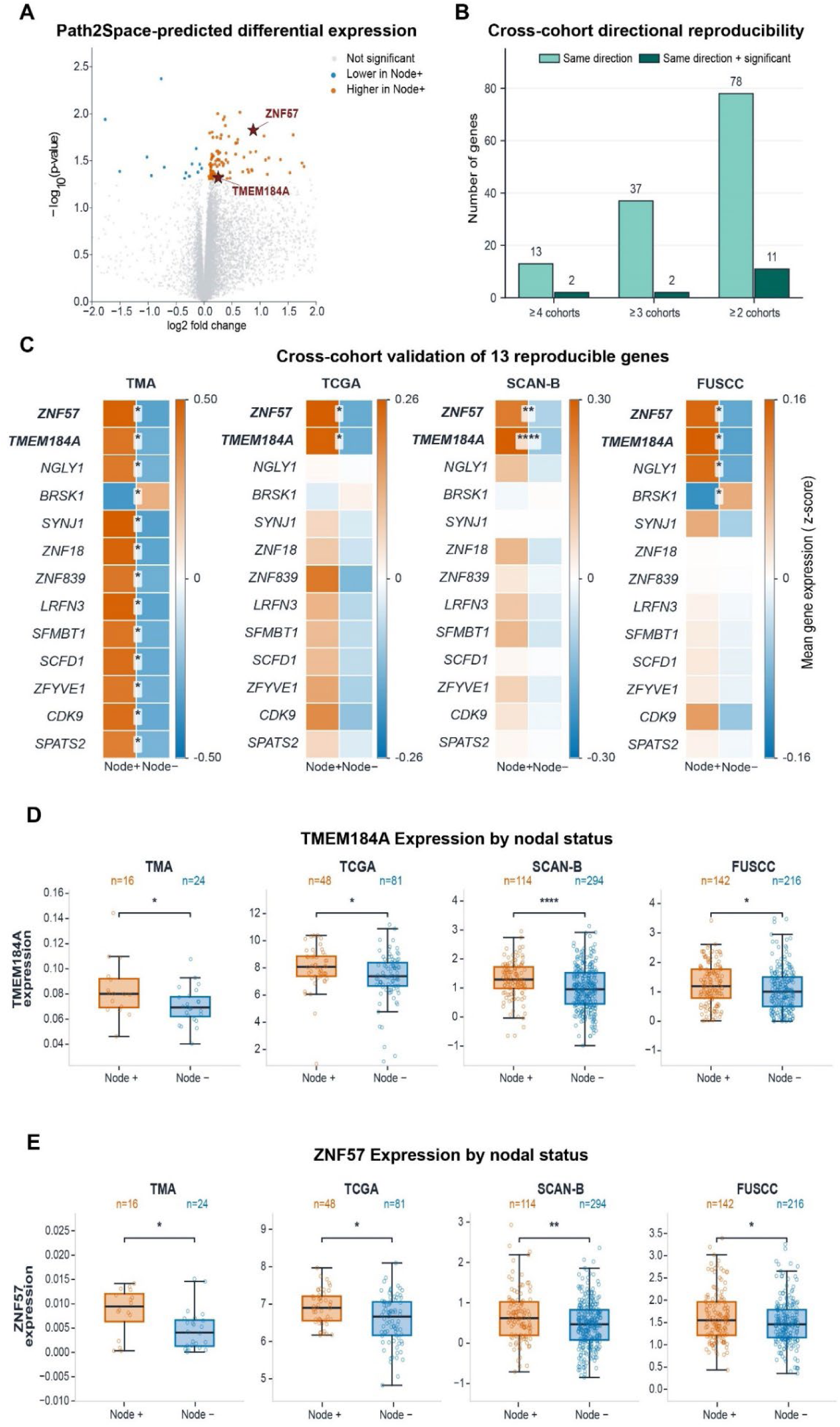
Cross-cohort validation identifies a conserved metastasis-associated gene signature centered on TMEM184A and ZNF57. **(A)** Differential expression analysis of Path2Space-predicted transcriptomes in the internal treatment-naïve TNBC discovery cohort comparing lymph node-positive (Node+) and lymph node-negative (Node-) primary tumors. The volcano plot highlights genes significantly associated with nodal metastasis. Differential expression was assessed using Welch’s t-test (P < 0.05). TMEM184A and ZNF57 emerged among the top metastasis-associated candidate genes. **(B)** Cross-cohort directional reproducibility of metastasis-associated genes across the internal discovery cohort and three independent TNBC cohorts (TCGA, SCAN-B, and FUSCC). Bars indicate the number of genes maintaining the same direction of association with lymph node status across at least two, three, or all four cohorts. Dark green bars denote genes that were both directionally consistent and statistically significant, illustrating progressive refinement from 118 candidate genes to a small set of reproducible biomarkers. **(C)** Cross-cohort validation of the 13 reproducibly associated genes. Heatmaps show cohort-specific mean gene expression (z-score) in lymph node-positive and lymph node-negative tumors across the internal TMA cohort, TCGA-BRCA, SCAN-B, and FUSCC. Warmer colors indicate higher relative expression and cooler colors indicate lower relative expression within each cohort. Orange and blue columns represent Node+ and Node-tumors, respectively. Asterisks indicate nominal statistical significance from two-sided Welch’s t-tests. TMEM184A and ZNF57 were consistently enriched in node-positive tumors across all four independent cohorts. **(D)** TMEM184A expression according to lymph node status across the internal discovery cohort and three external validation cohorts. Box plots display median, interquartile range (IQR), and individual patient measurements. Elevated TMEM184A expression was consistently associated with lymph node-positive tumors across all cohorts. **(E)** ZNF57 expression according to lymph node status across the internal discovery cohort and external validation cohorts. Consistent with TMEM184A, ZNF57 expression was reproducibly increased in node-positive tumors across geographically independent patient populations, further supporting the robustness and generalizability of the two-gene metastatic signature. Two-sided Welch’s t-tests were used; statistical significance is indicated as P < 0.05 (*), P < 0.01 (**), P < 0.001 (***), and P < 0.0001 (****).

We next examined the reproducibility of the entire metastatic gene set across independent patient populations. Although individual gene associations varied among cohorts, directional concordance was retained for a subset of genes **(Figure 2B)**. Overall, 78 genes retained the same direction of association with lymph node status in at least two cohorts, 37 were directionally concordant across three cohorts, and 13 preserved the same direction of association across all four datasets.

Further evaluation of these 13 directionally conserved genes revealed substantial variation in the magnitude and statistical strength of their associations across cohorts **(Figure 2C)**. Notably, ZNF57 and TMEM184A were the only genes that remained statistically significant in all four datasets, indetifying them as the two most reproducible metastasis-associated candidates.

Because TMEM184A and ZNF57 were the only genes that remained significantly associated with lymph node status across all four datasets, we next evaluated their individual expression across cohorts. TMEM184A was consistently elevated in lymph node-positive tumors in the discovery cohort and all three validation datasets **(Figure 2D)**. Likewise, ZNF57 demonstrated significantly higher expression in node-positive tumors across all four cohorts **(Figure 2E)**. The concordance of these associations between H&E-derived virtual transcriptomes and independently measured RNA-sequencing datasets further supports the reproducibility of TMEM184A and ZNF57 as metastasis-associated signals in TNBC.

### A two-gene metastatic score integrates complementary information from TMEM184A and ZNF57

Having identified TMEM184A and ZNF57 as the two genes that most consistently associated with lymph node metastasis across all four cohorts, we next asked whether combining these biomarkers could improve the ability to distinguish node-positive from node-negative tumors compared with either gene alone. Within each cohort, expression of TMEM184A and ZNF57 was independently standardized using gene-wise z-scores, and a fixed Virtual Metastatic Prediction Score (VMPS) was calculated as the average of the two standardized values **(Figure 3A)**. Because the VMPS contains only two predefined genes and requires neither model fitting nor cohort-specific optimization, it provides a simple and readily interpretable measure of metastatic transcriptional state.

**Figure 3.**
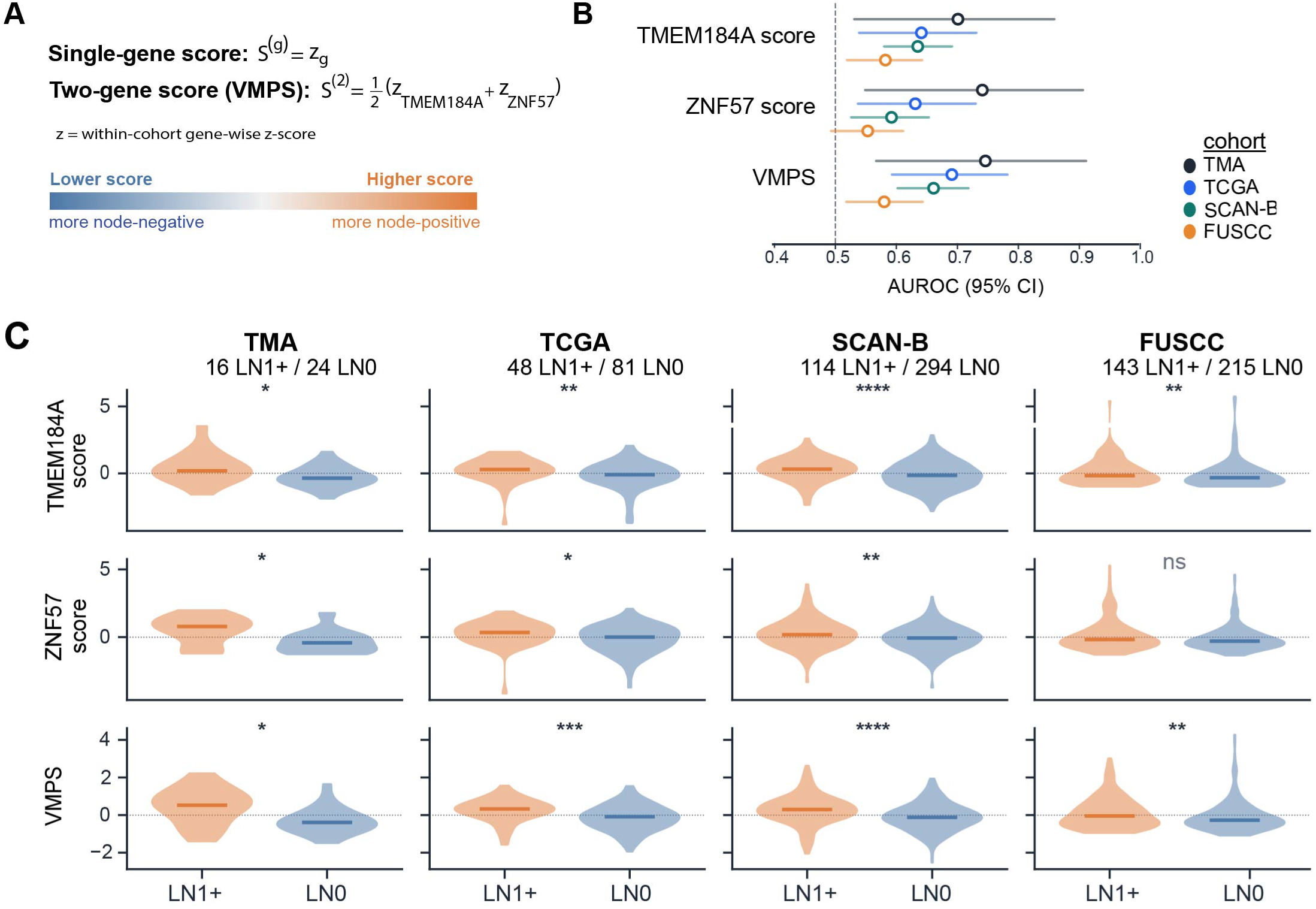
Development and cross-cohort assessment of a fixed two-gene Virtual Metastatic Prediction Score (VMPS) based on TMEM184A and ZNF57. **(A)** Construction of the Virtual Metastatic Prediction Score (VMPS). Gene expression values were standardized within each cohort using gene-wise z-scores. Individual gene scores correspond to the standardized expression of TMEM184A or ZNF57, whereas the composite two-gene score (VMPS) was calculated as the mean standardized expression of both genes. Higher scores indicate greater similarity to the node-positive transcriptional phenotype. **(B)** Comparison of discrimination for regional lymph-node involvement using TMEM184A alone, ZNF57 alone, or VMPS across the internal TMA cohort (n=40) and the TCGA-BRCA (n=129), SCAN-B (n=408), and FUSCC (n=358) external cohorts. Points show AUROC estimates and horizontal lines show 95% confidence intervals calculated using 2,000 patient-level bootstrap resamples. **(C)** Patient-level distributions of TMEM184A, ZNF57, and VMPS scores by regional lymph-node status in the curated internal TMA cohort and the TCGA-BRCA, SCAN-B, and FUSCC external cohorts. Violin plots show the distributions and individual patients; black horizontal bars and vertical lines indicate the group mean and 95% confidence interval. LN1+ and LN0 groups were compared using two-sided Mann–Whitney U tests; statistical significance is indicated as P < 0.05 (*), P < 0.01 (**), P < 0.001 (***), and P < 0.0001 (****); ns, not significant.

We next compared the performance of the individual gene scores and the VMPS across the discovery cohort and the three independent evaluation cohorts **(Figure 3B)**. Both TMEM184A and ZNF57 individually demonstrated reproducible discrimination of lymph node status, with AUROC values ranging from 0.58–0.70 and 0.55–0.72, respectively. The VMPS consistently achieved the highest or near-highest discriminatory performance across all cohorts, with AUROC values of 0.73 in the discovery cohort, 0.69 in TCGA-BRCA, 0.66 in SCAN-B, and 0.58 in FUSCC. Although the improvement over the strongest individual biomarker was modest, the reproducibility of the VMPS across independent populations suggests that TMEM184A and ZNF57 capture complementary rather than redundant aspects of metastatic biology.

We next examined VMPS distributions according to lymph node status in each cohort **(Figure 3C)**. TMEM184A scores were significantly elevated in node-positive tumors across all four datasets, whereas ZNF57 showed significant differences in the discovery cohort, TCGA-BRCA, and SCAN-B, with a similar but non-significant trend in FUSCC. Importantly, the VMPS consistently produced the clearest separation between node-positive and node-negative tumors, reaching statistical significance in every cohort despite substantial differences in patient populations, sequencing platforms, and cohort sizes.

Together, these findings demonstrate that the VMPS integrates complementary transcriptional information into a reproducible and interpretable quantitative measure associated with lymph node metastasis across independent patient cohorts.

### TMEM184A and ZNF57 define biologically distinct and robust transcriptional modules associated with metastatic progression

Although TMEM184A and ZNF57 emerged as the two most reproducible metastasis-associated genes across independent TNBC cohorts, individual biomarkers rarely function in isolation. Instead, they are expected to represent broader transcriptional programs that coordinate tumor behavior. To identify the biological programs represented by these biomarkers, we interrogated their local neighborhoods within a TNBC-specific co-expression network generated entirely from measured bulk and single-cell transcriptomic data.

The TMEM184A neighborhood formed a densely connected co-expression module containing representative membrane-associated proteins (SDCBP2, PROM2, TRPV6, CD82), glycosylation enzymes (FUT2 and FUT3), secreted proteins (CST6, PRSS22), and epithelial-associated genes including WNT7B and PSCA **(Figure 4A, Supplementary Table S3)**. Collectively, the composition of this module was consistent with an epithelial transcriptional program involving cell-surface organization, differentiation, and cell-cell communication. Functional enrichment analysis further supported this interpretation, revealing significant enrichment for endothelial cell migration, calcium-independent cell-cell adhesion, regulation of epidermal cell differentiation, acute-phase response, and inflammatory response to antigenic stimulus **(Figure 4B)**. These biological associations were further supported across multiple pathway annotation resources, including MSigDB Hallmark, Reactome, and WikiPathways, which identified epithelial remodeling, inflammatory signaling, secretory pathways, and TROP2-related signaling among the features associated with the TMEM184A module (**Supplementary Figure S4A**). Together, these findings indicate that TMEM184A marks a coherent epithelial remodeling program encompassing processes related to cell adhesion, differentiation, and tumor– microenvironment interactions.

**Figure 4.**
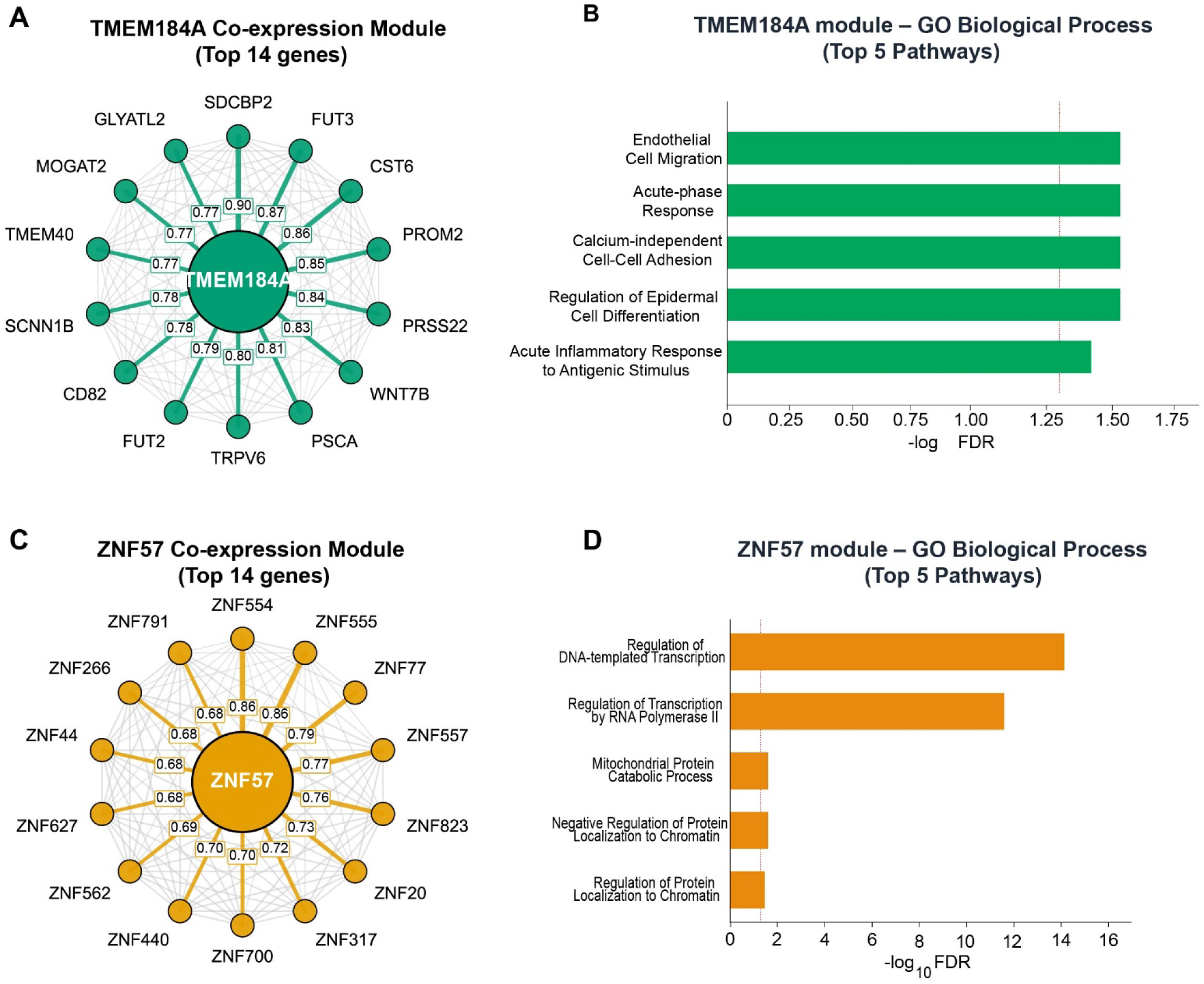
TMEM184A defines an epithelial differentiation and adhesion program, whereas ZNF57 defines a KRAB zinc-finger transcriptional regulatory program. **(A)** Co-expression network centered on TMEM184A within the TNBC-specific gene co-expression map. The network displays the 14 genes with the highest cosine similarity to TMEM184A among the 100-gene TMEM184A module. Representative genes include membrane associated proteins (PROM2, TRPV6, and CD82), glycosylation enzymes (FUT2 and FUT3), and secreted proteins (CST6 and PRSS22), suggesting coordinated regulation of epithelial differentiation, cell adhesion, and cell-surface biology. Numbers on the connecting edges indicate pairwise cosine similarity between connected genes. **(B)** Gene Ontology (GO) biological process enrichment analysis of the 100-gene TMEM184A co-expression module. The top five significantly enriched GO biological processes (FDR < 0.05) are shown, ranked by −log10 (FDR). The TMEM184A module was enriched for biological processes related to endothelial and epithelial biology, cell migration, cell adhesion, inflammatory responses, and regulation of epithelial cell differentiation, supporting its association with epithelial plasticity and metastatic progression. **(C)** Co-expression network centered on ZNF57 within the TNBC-specific gene co-expression map. The network displays the 14 genes with the highest cosine similarity to ZNF57 among the 52-gene ZNF57 module, which is composed predominantly of KRAB zinc-finger transcription factors. Numbers on the hub-to-gene spokes indicate cosine similarity between each displayed gene and its TMEM184A or ZNF57 hub. **(D)** Gene Ontology biological process enrichment analysis of the 52-gene ZNF57 module. The top five significantly enriched GO biological processes (FDR < 0.05) are shown, ranked by −log10 (FDR). The ZNF57 module was enriched for biological processes related to transcriptional regulation, chromatin-associated protein localization, and mitochondrial protein catabolism, consistent with a transcriptional regulatory program distinct from the TMEM184A-associated epithelial adhesion program.

In contrast, the ZNF57 neighborhood formed a distinct co-expression module composed predominantly of KRAB zinc-finger transcription factors, including ZNF554, ZNF555, ZNF77, ZNF557, ZNF823, and related family members **(Figure 4C, Supplementary Table S4)**. Collectively, the composition of this module was consistent with a coordinated transcriptional regulatory program distinct from the epithelial remodeling program represented by the TMEM184A module. Functional enrichment analysis further supported this interpretation, revealing significant enrichment for DNA-templated transcription, RNA polymerase II-mediated transcription, regulation of protein localization to chromatin, and mitochondrial protein catabolic processes **(Figure 4D)**. Complementary pathway databases further supported this interpretation, identifying RNA polymerase II transcription, transcriptional regulation, and gene-expression pathways among the principal biological functions associated with the ZNF57 (**Supplementary Figure S4B**). Together, these findings indicate that the ZNF57 marks a transcriptional regulatory program centered on transcriptional and chromatin-associated processes.

Because both modules were derived computationally from the same co-expression landscape, we next evaluated whether they represented independent biological programs rather than different manifestations of a common epithelial transcriptional state. Across three independent TNBC single-cell RNA-sequencing atlases, TMEM184A and ZNF57 module activities exhibited consistently lower correlations than expected from size-matched random gene sets, demonstrating that the two modules capture complementary rather than redundant transcriptional programs within cancer epithelial cells (**Supplementary Figure S4C**). Thus, although both modules are associated with metastatic progression, they appear to represent distinct biological dimensions of the metastatic phenotype.

We next asked whether the identified modules were robust to alternative network construction strategies. Reconstruction of the TNBC co-expression map using bulk RNA-sequencing data alone, completely excluding single-cell information, recovered both TMEM184A and ZNF57 modules with substantial overlap relative to the full network, greatly exceeding chance expectation (**Supplementary Figure S4D**). Furthermore, both modules exhibited significantly greater within-module co-expression than size-matched random gene sets across the independent TCGA, SCAN-B, and FUSCC cohorts (**Supplementary Figure S4E**), confirming that these modules represent coherent and reproducible transcriptional programs rather than arbitrary collections of correlated genes.

Finally, because robust biomarkers should remain stable despite technical variability, we assessed the effect of progressively increasing measurement noise on both module scores and individual hub-gene expression. Module-based scores consistently retained substantially higher agreement with their original values than expression of TMEM184A or ZNF57 alone (**Supplementary Figure S4F**), demonstrating that aggregating coordinated transcriptional information substantially improves analytical robustness relative to single-gene measurements.

Collectively, these analyses demonstrate that TMEM184A and ZNF57 define biologically distinct yet complementary transcriptional modules within TNBC. The TMEM184A module captures an epithelial remodeling program involving cell-surface organization, adhesion, differentiation, and tumor– microenvironment interactions, whereas the ZNF57 module captures a KRAB zinc-finger-centered transcriptional regulatory program with chromatin-associated features. Together, their biological coherence, reproducibility, and independence support the existence of two distinct transcriptional programs associated with lymp node metastasis in TNBC.

### TMEM184A and ZNF57 modules define complementary molecular states across TNBC subtypes

Having established that TMEM184A and ZNF57 define biologically distinct transcriptional programs, we next asked how these programs relate to the molecular heterogeneity of TNBC. We examined module activity across two-independent subtype classification systems using the TCGA and FUSCC cohorts.

In the TCGA cohort, classified according to the refined TNBCtype-4 framework (BL1, BL2, M, and LAR), the TMEM184A module exhibited pronounced subtype-specific differences, with higher activity in LAR and BL2 tumors and lower activity in BL1 and M tumors (**Figure 5A**). Overall subtype differences were highly significant (Kruskal-Wallis, *P* = 2.5 × 10⁻⁵), In contrast, the ZNF57 module showed no significant subtype-specific variation (*P* = 0.232), whereas VMPS differed significantly across subtypes (*P* = 6.5 × 10⁻⁴).

**Figure 5.**
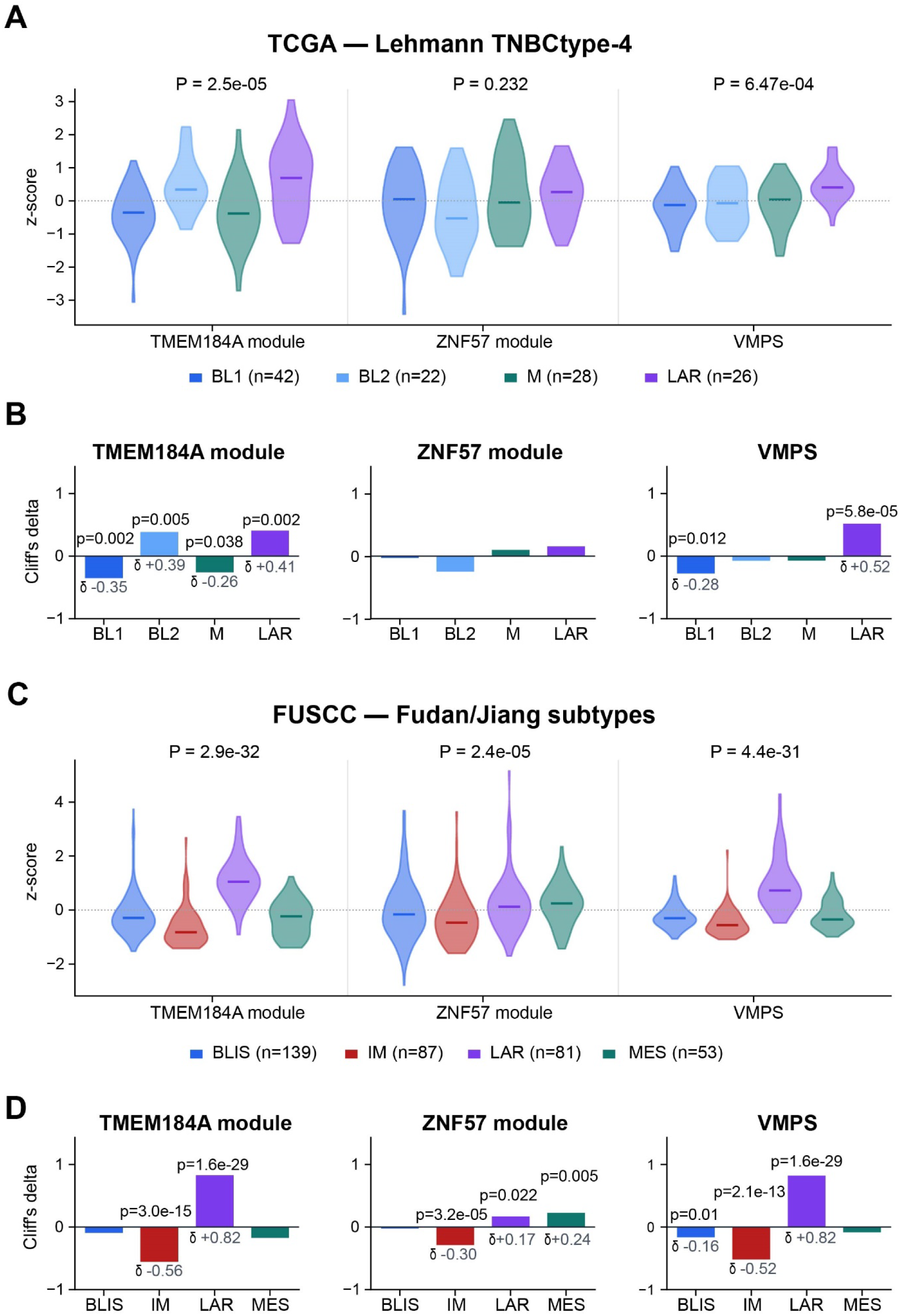
TMEM184A and ZNF57 transcriptional programs exhibit distinct relationships with TNBC molecular subtypes across independent classification systems. **(A)** Distribution of TMEM184A-module, ZNF57-module, and VMPS scores across the four refined Lehmann TNBCtype-4 subtypes in TCGA: basal-like 1 (BL1), basal-like 2 (BL2), mesenchymal (M), and luminal androgen receptor (LAR). Violin plots show standardized scores among 118 tumors with published TNBCtype-4 calls (BL1, n = 42; BL2, n = 22; M, n = 28; LAR, n = 26). The TCGA subtype assignments were taken from Lehmann et al.^5^ Supplementary Table S3 and were not supplied natively by TCGA. Scores differed across TNBCtype-4 subtypes for the TMEM184A module (Kruskal–Wallis H = 24.03, P = 2.46×10⁻⁵) and VMPS (H = 17.19, P = 6.47×10⁻⁴), but not for the ZNF57 module (H = 4.29, P = 0.232). **(B)** One-versus-rest effect-size analysis in TCGA. Bars show Cliff’s delta (δ) for each TNBCtype-4 subtype relative to all other TCGA subtypes combined; positive values indicate higher scores and negative values indicate lower scores. Two-sided Mann–Whitney U tests were used for nominal P values. Subtypes with significant p values are only shown. The TMEM184A module and VMPS showed their strongest positive effects in LAR, whereas the ZNF57 module showed weaker and nonsignificant subtype-specific effects. **(C)** Validation in the independent FUSCC cohort using the native four-class Fudan/Jiang TNBC subtype system: basal-like immunosuppressed (BLIS), immunomodulatory (IM), mesenchymal (MES), and luminal androgen receptor (LAR). Violin plots show standardized scores among 360 tumors with subtype and nodal-status data. Scores differed across subtypes for the TMEM184A module (Kruskal– Wallis H = 149.79, P = 2.92×10⁻³²), ZNF57 module (H = 24.11, P = 2.38×10⁻⁵), and VMPS (H = 144.32, P = 4.43×10⁻³¹). The Fudan/Jiang and Lehmann subtype systems are shown as cohort-specific classifications and are not treated as interchangeable. **(D)** One-versus-rest effect-size analysis in FUSCC. Bars show Cliff’s delta (δ) for each native Fudan/Jiang subtype relative to all other FUSCC subtypes combined. Two-sided Mann–Whitney U tests were used for nominal P values. Subtypes with significant p values are only shown. TMEM184A-module, ZNF57-module, and VMPS scores showed positive effects in LAR and negative effects in IM; the ZNF57-module pattern was weaker than the TMEM184A-module and VMPS patterns.

One-versus-rest effect-size analysis further resolved these subtype-associated patterns (**Figure 5B**). The TMEM184A module was enriched in LAR (Cliff’s δ = 0.41) and in BL2 (δ = 0.39) tumors and depleted in BL1 (δ = −0.35) and M tumors (δ = −0.26). In contrast, the ZNF57 module exhibited uniformly small effect sizes and no significant subtype enrichment. VMPS showed its strongest association with the LAR subtype (δ = 0.52), with modest depletion in BL1 tumors (δ = −0.28) and minimal effects in BL2 and M tumors. Thus, although both the TMEM184A module and VMPS were associated with TNBC subtype, their patterns were distinct: the TMEM184A module captured borader subtype-associated variation, whereas VMPS was predominantly associated with LAR and BL1 subtype.

We next asked whether these subtype-associated patterns generalized to an independent patient population using the FUSCC cohort, classified according to the **Fudan/Jiang** TNBC taxonomy (BLIS, IM, MES, and LAR). Consistent with the TCGA findings, both TMEM184A module activity and VMPS demonstrated highly significant subtype-specific differences (both *P* < 10⁻⁵), with maximal activity observed in LAR tumors (**Figure 5C**). Unlike TCGA, however, the ZNF57 module also showed significant subtype-associated variation (*P* = 2.4 × 10⁻⁵), indicating a more context-dependent relationship with TNBC molecular subtype.

Effect-size analysis further resolved these patterns **(Figure 5D)**. The TMEM184A module showed strong enrichment in LAR tumors (δ = 0.82) and depletion in IM (δ = −0.56) tumors. The ZNF57 module demonstrated modest but significant enrichment in LAR (δ = 0.17) and MES tumors (δ = 0.24), together with depletion in IM tumors (δ = −0.30). VMPS showed strong enrichment in LAR tumors (δ = 0.82) and marked depletion in IM tumors (δ = −0.52), with more modest depletion in BLIS tumors (δ = −0.16). Across the two classification systems, the most reproducible feature was enrichment of the TMEM184A module and VMPS in LAR tumors. In contrast, ZNF57 showed little subtype specificity in TCGA but modest subtype-associated variation in FUSCC, indicating a more context-dependent relationship with TNBC molecular taxonomy.

Given the reproducible enrichment of the TMEM184A module and VMPS in LAR tumors, we next quantified how strongly the individual genes, their corresponding transcriptional modules, and VMPS discriminated each TNBC subtype from all other tumors (**Supplementary Figure S5A,B**). Using one-versus-rest AUROC analysis, TMEM184A expression, TMEM184A module activity, and VMPS showed the strongest discrimination of LAR tumors in both FUSCC and TCGA. In FUSCC, these features also showed AUROC values below 0.5 for IM tumors, indicating an inverse association with the IM subtype. In contrast, ZNF57 expression and module activity showed weaker and less consistent subtype discrimination across cohorts.

Because these discrimination patterns suggested a prominent relationship with LAR biology, we next examined the association of each feature with an independently defined luminal/androgen receptor transcriptional axis. TMEM184A expression, TMEM184A module activity, and VMPS showed the strongest positive correlations with this axis in both cohorts, whereas ZNF57 expression and module activity showed weaker and less consistent associations (**Supplementary Figure S5C**). These findings indicate that the reproducible LAR enrichment of the TMEM184A program reflects its association with a broader luminal/androgen receptor transcriptional state rather than simply a categorical subtype label.

Together, these analyses reveal distinct relationships between the metastasis-associated transcriptional programs and established TNBC molecular heterogeneity. The TMEM184A module showed the strongest and most reproducible relationship with subtype biology, particularly the luminal/androgen receptor state, whereas ZNF57 displayed weaker and more context-dependent subtype specificity. VMPS was likewise enriched in LAR tumors but exhibited a subtype pattern distinct from the broader TMEM184A module, supporting their complementary roles as a biological transcriptional program and a metastasis-associated two-gene score, respectively.

### Virtual transcriptomic biomarkers provide robust cross-cohort metastatic risk stratification

Having characterized the biological programs represented by these virtual transcriptomic biomarkers, we next evaluated whether they provide reproducible measures of metastatic risk across independent patient populations. We therefore examined whether increasing biomarker scores were consistently associated with lymph node metastasis across independent TNBC cohorts.

We first quantified the associations of module scores and VMPS with nodal involvement using logistic regression in the TCGA, SCAN-B, and FUSCC cohorts **(Figure 6A)**. The TMEM184A module demonstrated consistently positive effect estimates across all three datasets, with odds ratios (OR) per 1-standard deviation (SD) ranging from 1.35 to 1.55. Meta-analysis across cohorts yielded a pooled OR of 1.44 (95% CI 1.25–1.67) with no observed between-cohort heterogeneity (I² = 0%), supporting a reproducible association between the TMEM184A transcriptional and lymph node metastasis. In contrast, the ZNF57 module exhibited greater between-cohort variability, with a significant positive association in SCAN-B, a positive but nonsignificant association in TCGA, and no association in FUSCC (pooled OR 1.24, I² = 80%). VMPS showed consistent positive associations across all three cohorts, yielding a pooled odds ratio of 1.58 (95% CI 1.28–1.94) with moderate heterogeneity (I² = 44%). Complementary AUROC analyses further evaluated the ability of the individual genes, VMPS, and module-based scores to distinguish node-positive from node-negative tumors across cohorts (**Supplementary Figure S6A**). Adjustment for estimated tumor purity or proliferation had minimal effect on these associations, suggesting that they were not explained simply by differences in tumor cellularity or proliferative activity (**Supplementary Figure S6B**).

**Figure 6.**
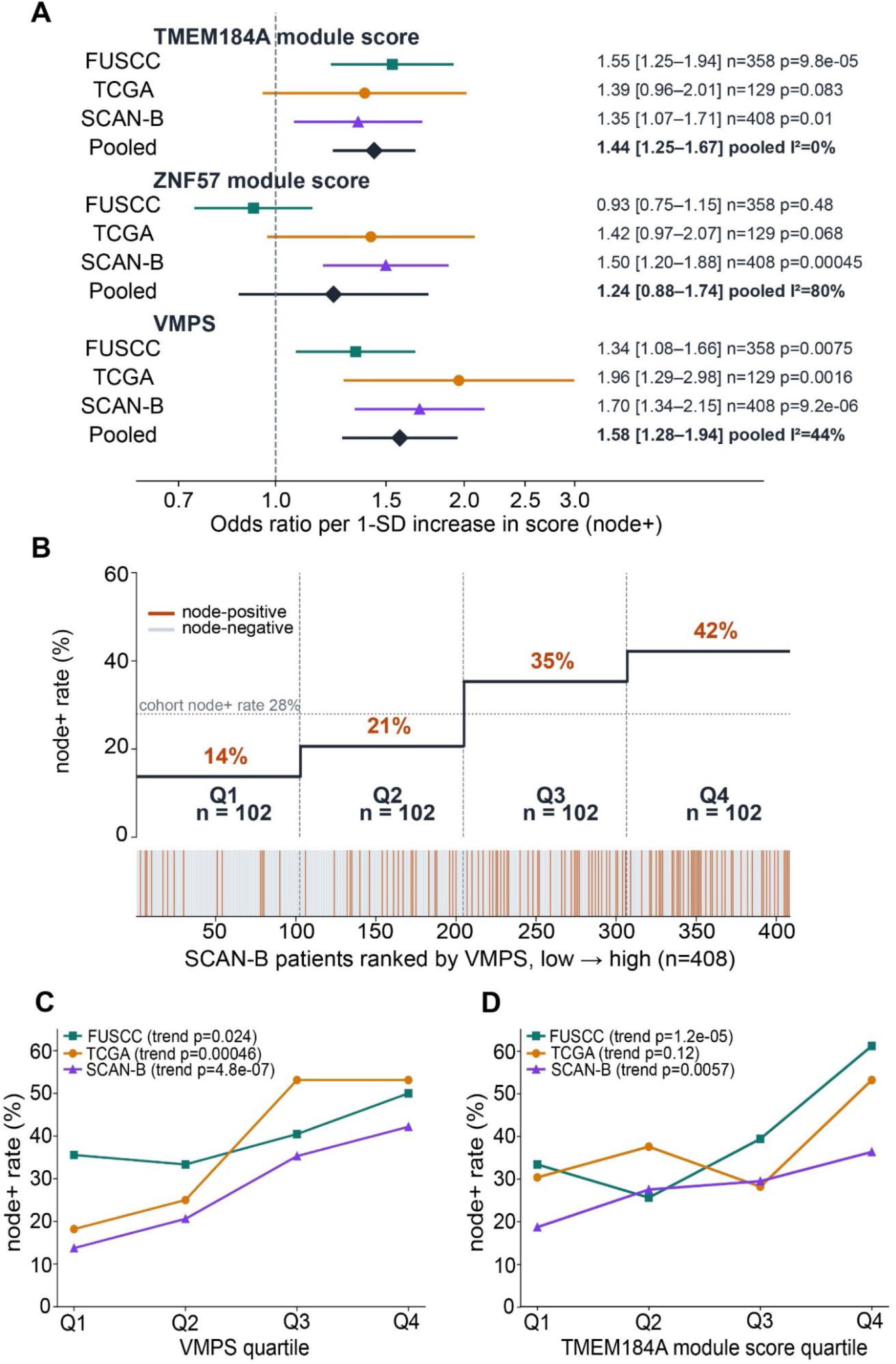
Virtual transcriptomic programs define reproducible gradients of lymph-node involvement across independent TNBC cohorts. **(A)** Cross-cohort meta-analysis of associations between virtual transcriptomic scores and lymph node positivity. Forest plots show odds ratios (ORs) with 95% confidence intervals for a one-standard-deviation increase in the TMEM184A module score, ZNF57 module score, and VMPS across the FUSCC, TCGA-BRCA, and SCAN-B cohorts. Random-effects pooled estimates and between-study heterogeneity (I²) are shown. The TMEM184A module and VMPS demonstrated consistent positive associations with regional lymph node metastasis across cohorts, whereas the ZNF57 module exhibited greater between-cohort heterogeneity. **(B)** Patient-level risk stratification using VMPS in the independent SCAN-B cohort. Patients were ranked according to the VMPS and divided into quartiles (Q1–Q4). The observed prevalence of lymph node-positive disease increased progressively from 14% in Q1 to 42% in Q4, exceeding the overall cohort prevalence of 28%. The raster plot below illustrates the distribution of node-positive and node-negative patients across the ranked cohort. This panel illustrates the quartile-based stratification framework used for analyses in Panels C and D. **(C)** Cross-cohort assessment of VMPS quartiles and regional lymph-node involvement. Patients in FUSCC, TCGA-BRCA, and SCAN-B were grouped into cohort-internal VMPS quartiles. Rates of lymph node positivity showed positive ordered trends across quartiles in all three cohorts, although adjacent quartiles were not uniformly monotonic in every cohort. Nominal P values were calculated using two-sided Cochran-Armitage tests for trend across quartiles coded 1-4. **(D)** Cross-cohort assessment of TMEM184A-module quartiles and regional lymph-node involvement. Patients in FUSCC, TCGA-BRCA, and SCAN-B were grouped into cohort-internal TMEM184A-module-score quartiles. Rates of lymph node positivity showed positive ordered trends in FUSCC and SCAN-B, although adjacent quartiles were not uniformly monotonic. Nominal P values were calculated using two-sided Cochran-Armitage tests for trend across quartiles coded 1-4.

To determine whether the VMPS provides clinically meaningful risk stratification, we ranked all SCAN-B tumors according to VMPS and divided patients into equal-risk quartiles. The prevalence of lymph node metastasis increased progressively from **14% in the lowest-risk quartile (Q1)** to **42% in the highest-risk quartile (Q4) (Figure 6B).** This monotonic increase is consistent with a graded association between VMPS and regional lymph-node involvement. Across SCAN-B, TCGA, and FUSCC, VMPS quartiles showed positive ordered trends in lymph node positivity, although adjacent quartiles were not uniformly monotonic in every cohort **(Figure 6C)**. Positive ordinal-quartile trends were observed in all three cohorts despite differences in patient populations and sequencing platforms, supporting a reproducible association between higher VMPS and regional lymph-node involvement across independent datasets.

Finally, we performed the same analysis using the TMEM184A module score **(Figure 6D)**. TMEM184A module-score quartiles showed significant positive ordered trends in nodal involvement in SCAN-B and FUSCC, although adjacent quartiles were not uniformly monotonic. A similar overall pattern was observed in TCGA, although the trend did not reach statistical significance. The TMEM184A module also retained discriminatory performance across multiple molecular subtype contexts, although estimates were less precise in smaller subgroup analyses (**Supplementary Figure S6C**). In explaratory models incorporating clinicopathologic variables, addition of either the TMEM184A module or VMPS showed variable improvements in cross-validated discrimination across cohorts (**Supplementary Figure S6D**). These findings suggest that both VMPS and TMEM184A module may provide information beyond conventional clinicopathologic variables, without establishing superiority of one over the other.

Together, these findings demonstrate that VMPS and the TMEM184A module provide complementary measures of metastasis-associated biology in TNBC. VMPS defines a reproducible gradient of lymph node involvement across independent cohorts, whereas the TMEM184A module captures a broader transcriptional program with highly consistent cross-cohort associations. These findings support the potential of both gene- and module-based virtual transcriptomic features for metastatic risk stratification in TNBC.

## DISCUSSION

### Virtual transcriptomics enables scalable discovery of metastatic programs from routine histopathology

Routine histopathology remains the cornerstone of breast cancer diagnosis, yet its interpretation has traditionally been confined to morphologic assessment. Recent advances in computational pathology have demonstrated that deep learning models can infer genomic alterations, molecular subtypes, immune states, and clinical outcomes directly from histologic architecture^17,35–37^, suggesting that tissue morphology encodes rich molecular information. Here, we extend this paradigm by using virtual transcriptomics to bridge digital pathology with molecular discovery. By reconstructing genome-wide transcriptional profiles from H&E images using Path2Space, we interrogated the inferred molecular landscape to identify conserved metastatis-associated transcriptional programs and their underlying biology.

A key conceptual advance of this study is the emphasis on biological reproducibility rather than discovery alone. Starting from an internal treatment-naïve TNBC cohort, we systematically validated candidate genes across three independent international RNA-sequencing cohorts representing distinct patient populations, sequencing platforms, and geographic regions. The progressive reduction from 118 candidate genes to only two reproducible markers illustrates how virtual transcriptomics, when coupled with rigorous cross-cohort validation, can identify metastatic programs that generalize across independent datasets rather than reflecting cohort-specific transcriptional variation. This workflow establishes a scalable framework for converting routine pathology slides into biologically interpretable molecular hypotheses without requiring RNA sequencing.

### A parsimonious two-gene signature captures a conserved metastatic transcriptional state

A central finding of this study is that only TMEM184A and ZNF57 remained consistently associated with nodal metastasis across all four cohorts. Although numerous genes exhibited differential expression within individual datasets and a subset retained directional concordance across cohorts, no other genes were significantly associated with lymph node status in all four datasets. This convergence suggests that metastatic biology in TNBC may be represented by relatively stable transcriptional programs despite extensive genomic heterogeneity.

Importantly, VMPS generalized more effectively than larger candidate signatures. Increasing the number of genes modestly improved performance within the discovery cohort but failed to improve - and in some cases reduced - performance in external validation datasets. Such findings reinforce an increasingly recognized principle in biomarker development: parsimonious models frequently exhibit greater transportability than highly complex signatures that capture cohort-specific transcriptional variation.

Equally important, TMEM184A and ZNF57 should not be viewed as isolated biomarkers but as representative nodes within complementary transcriptional networks. Their association with epithelial remodeling and transcriptional regulatory programs, respectively, likely explains why a simple two-gene score generalized robustly across independent cohorts despite the molecular heterogeneity of TNBC.

### TMEM184A identifies an epithelial remodeling program linked to metastatic competence

TMEM184A emerged as the central node of a highly coherent transcriptional module characterized by genes involved in cell-surface organization, glycosylation, and epithelial biology. Functional enrichment further linked this module to cell-cell adhesion, epithelial differentiation, inflammatory signaling, and epithelial remodeling. Collectively, these processes are consistent with a transcriptional state that may influence interactions between tumor cells and their surrounding microenvironment during metastatic progression.

The coexistence of epithelial features with pathways related to epithelial remodeling is consistent with the emerging concept of hybrid epithelial states^38^, in which tumor cells retain epithelial characteristics while acquiring features associated with cellular plasticity and metastatic behaviour. The presence of genes involved in cell-surface organization, glycosylation, and epithelial communication, together with enrichment of pathways related to cell-cell adhesion, epithelial differentiation, inflammatory signaling, and epithelial remodeling, suggests that this transcriptional program may facilitate dynamic interactions between tumor cells and the surrounding microenvironment during metastatic progression.

Although TMEM184A has received relatively little attention in breast cancer biology, our findings identify it as a previously underappreciated candidate regulator of metastatic competence that warrants further future mechanistic investigation. As a membrane-associated protein, TMEM184A may also represent an attractive candidate for future therapeutic targeting.

### ZNF57 identifies a complementary transcriptional regulatory program

In contrast to epithelial remodeling program associated with TMEM184A, ZNF57 emerged as the central node of a distinct transcriptional module dominated by KRAB zinc-finger proteins and pathways related to transcriptional and chromatin regulation. KRAB zinc-finger proteins comprise a large family of transcriptional regulators that can recruit chromatin-modifying complexes and establish transcriptionally repressive states^39^, suggesting that the ZNF57 module may capture a regulatory dimension of metastasis-associated biology distinct from the cell-surface and microenvironmental processes associated with TMEM184A.

Although the broader ZNF57 module exhibited greater variability across independent cohorts than the TMEM184A module, ZNF57 itself consistently emerged as one of only two reproducible metastatis-associated genes across all four datasets. This observation suggests that ZNF57 may serve as a robust marker of a conserved transcriptional regulatory state even when the broader regulatory network varies across patient populations. Together, TMEM184A and ZNF57 therefore capture complementary dimensions of metastatic biology: one characterized by epithelial remodeling and tumor-microenvironment interactions and the other by transcriptional and chromatin-associated regulation.

### Histology-derived virtual transcriptomics enables clinically interpretable metastatic risk stratification

From a translational perspective, an important objective of computational pathology is not simply accurate prediction, but clinically meaningful patient stratification. The present study demonstrates that both the TMEM184A module and the parsimonious two-gene VMPS generate reproducible and clinically interpretable gradients of metastatic risk across three independent international cohorts.

Rather than functioning as binary classifiers, higher VMPS and TMEM184A module activity showed positive ordered associations with rates of regional lymph node metastasis, indicating that these virtual transcriptomic features capture continuous biological variation in metastatic competence. The consistency of these trends across TCGA, SCAN-B, and FUSCC supports their potential generalizability and suggests that routine histopathology may provide clinically relevant molecular information without additional tissue processing.

Importantly, the virtual transcriptomic biomarkers also showed reproducible associations with established TNBC molecular subtype classifications. TMEM184A and VMPS were consistently enriched in the LAR subtype across independent classification systems, linking these biomarkers to a recognized biological subtype. These observations suggest that virtual transcriptomics complements existing molecular classifications by providing biologically interpretable information on metastasis-associated transcriptional programs within established TNBC subtypes.

Because the approach operates on routinely acquired H&E slides, it provides a potential route toward integration with existing digital pathology workflows without requiring additional tissue acquisition or molecular assays. This makes virtual transcriptomics particularly well suited for large retrospective clinical cohorts and settings where comprehensive transciptomic profiling is unavailable.

### Virtual transcriptomics bridges digital pathology and spatial molecular biology

Our study contributes to the rapidly evolving convergence of digital pathology and spatial molecular biology. Spatial transcriptomic technologies provide unprecedented molecular resolution but remain costly, technically demanding and difficult to implement routinely across large clinical cohorts^40,41^. Conversely, pathology foundation models have demonstrated remarkable ability to infer diverse molecular phenotypes directly from tissue morphology^36,42,43^, creating new opportunities to extract biological information from routinely acquired H&E images.

By combining foundation-model-derived virtual transcriptomics with transcriptional module analysis, our framework bridges these complementary approaches. Rather than replacing experimentally measured transcriptomics, virtual transcriptomics provides a scalable strategy for generating biologically interpretable molecular hypotheses that can subsequently be validated using spatial transcriptomics and functional studies.

### Study limitations

First, transcriptomic profiles in the discovery cohort were computationally inferred rather than experimentally measured, and the initial discovery cohort was relatively modest in size. Although extensive validation across three independent RNA-sequencing cohorts supports the biological relevance and reproducibility of the identified biomarkers, future studies integrating matched H&E images, bulk RNA sequencing, and spatial transcriptomics will be important for directly evaluating the molecular fidelity of virtual transcriptomics. Larger independent cohorts will subsequently be needed to establish its clinical utility.

Second, although our findings place TMEM184A and ZNF57 within biologically coherent transcriptional programs, their mechanistic contributions to metastatic progression remain to be established experimentally. Future functional studies will be required to determine whether these genes actively contribute to metastatic progression or primarily serve as biomarkers of broader transcriptional programs.

Finally, the present study focused on lymph node metastasis as a clinically relevant indicator of metastatic competence. Future studies should determine whether the identified virtual transcriptomic programs also predict distant metastasis, recurrence, treatment response, and other clinically-relevant outcomes.

## Conclusions

In summary, we demonstrate that AI-enabled virtual transcriptomics can transform routine histopathology into a scalable platform for molecular discovery in TNBC. By integrating digital pathology with foundation-model-based virtual transcriptomics and rigorous cross-cohort validation, we identified a parsimonious two-gene VMPS composed of TMEM184A and ZNF57 that reproducibly associates with lymph node metastasis across independent international cohorts.

Beyond identifying reproducible biomarkers, our study demonstrates that virtual transcriptomics provides biological context by placing candidate genes within complementary transcriptional programs linked to epithelial remodeling and transcriptional regulation. Collectively, these findings establish routine H&E histopathology as a source of biologically interpretable molecular information, enabling biomarker discovery without requiring experimental transcriptomic profiling.

## RESOURCE AVAILABILITY

### Lead contact

Further information and requests for resources and data should be directed to and will be fulfilled by the lead contacts, Mihriban Karaayvaz and Murat Yildirim.

### Materials availability

This study did not generate new unique reagents.

### Data and code availability

All original code will be deposited at Zenodo Digital repository as well as our lab’s github page (https://github.com/yyildirimlab) as of the date of publication. Any additional information required to reanalyze the data reported in this paper is available from the lead contact upon request.

### Ethics

The requirement for informed consent was waived by the Institutional Review Board (protocol 19-1257) because this was a minimal-risk study using data collected as part of routine clinical practice.

## ACKNOWLEDGEMENTS

The authors thank the Cleveland Clinic Research Image Core facility for histological staining services and the Department of Pathology archives for providing breast cancer specimens. The authors also thank Resul Ozbilgic for helpful discussions and technical input related to this work. This work was supported by US National Institute of Health (NIH) grants #R00EB027706 (to M.Y.), American Brain Tumor Association Discovery grant #DG2500080 in memory of Kaitlyn Berg (to M.Y.), Cleveland Clinic Research Startup (to M.Y.), Cleveland Clinic Research Startup (MK), Ohio Cancer Research (MK), R01CA257502 (RAK). This work was also supported by the National Cancer Institute of the National Institutes of Health through Case Comprehensive Cancer Center Support grant (P30CA043703).

## AUTHOR CONTRIBUTIONS

Conceptualization, A.S., M.K., and M.Y.; methodology, A.S., M.K., and M.Y.; software, A.S.; validation, A.S.; formal analysis, A.S.; investigation, A.S., M.K., and M.Y.; resources, M.K., M.N., D.D.S., R.A.K., and M.Y.; data curation, A.S.; writing - original draft, A.S., M.K., and M.Y.; writing - review and editing, A.S., M.K., and M.Y.; visualization, A.S., M.K., and M.Y.; supervision, M.K. and M.Y.; project administration, M.K. and M.Y.; funding acquisition, M.K., R.A.K., and M.Y.

## KEY RESOURCES TABLE

| **RESOURCE** | **SOURCE** | **IDENTIFIER** |
|---|---|---|
| Path2Space inference framework | Shulman et al.<sup>30</sup> | doi:10.1016/j.cell.2026.04.023 |
| Karaayvaz TNBC single-cell atlas | Karaayvaz et al.<sup>32</sup> | doi:10.1038/s41467-018-06052-0 |
| Wu breast-cancer single-cell atlas | Wu et al.<sup>33</sup> | GEO: GSE176078 |
| Pal breast single-cell atlas | Pal et al.<sup>34</sup> | doi:10.15252/embj.2020107333 |
| TNBCtype-4 classification | Lehmann et al.<sup>5</sup> | doi:10.1371/journal.pone.0157368 |
| FUSCC TNBC resource | Xiao et al.<sup>44</sup> | doi:10.1158/1078-0432.CCR-18-3524 |
| TCGA breast-cancer resource | TCGA Network<sup>45</sup> | doi:10.1038/nature11412 |
| CTransPath architecture | Wang et al.<sup>31</sup> | doi:10.1016/j.media.2022.102559 |
| gseapy | Fang et al.<sup>46</sup> | doi:10.1093/bioinformatics/btac757 |
| Enrichr | Kuleshov et al.<sup>47</sup> | doi:10.1093/nar/gkw377 |

## EXPERIMENTAL MODEL AND SUBJECT DETAILS

The study combined an internal TNBC TMA cohort with independent bulk transcriptomic cohorts and three single-cell breast-cancer atlases used for graph construction. The primary internal unit of analysis was the patient. Where a patient contributed more than one tissue core, inferential analyses used one patient-level observation; core-level summaries were exploratory. Regional nodal involvement (LN0 versus LN1+) was the primary outcome for the reported association analyses and was kept distinct from distant-metastasis and survival endpoints. The study cohorts, data types, samples sizes, and their respective analytical roles are summarized in **Table 1**. LN0 versus LN1+ denotes absence versus presence of regional lymph-node involvement; it is not a distant-metastasis endpoint.

**Table 1.** Study resources, analytical roles, and verified counts.

| Resource | Role | Verified analysis count | Outcome or annotation |
|---|---|---|---|
| Internal TMA | Discovery / image-expression analysis | 40 curated patient-level gene-analysis set | Regional nodal involvement: 16 LN1+ and 24 LN0 in curated set |
| TCGA TNBC | External nodal analysis | 129 (48 LN1+, 81 LN0) | Pathologic N0 versus N1–N3 |
| SCAN-B | External nodal analysis | 408 (114 LN1+, 294 LN0) | NodeNegative versus NodePositive |
| FUSCC | External nodal analysis | 358 (143 LN1+, 215 LN0) | Observed LN_positive only |
| FUSCC | Subtype-only analysis | 360 | Fudan/Jiang subtype labels |
| Single-cell atlases | TNBC graph evidence | Karaayvaz et al.<sup>32</sup>, Wu et. al<sup>33</sup>, Pal et. al<sup>34</sup> | Epithelial-cell expression correlations |

## METHOD DETAILS

### Internal TMA preparation, imaging, and curation

42 patients were technically available to the internal image-expression workflow (17 LN1+ and 25 LN0). The primary curated patient-level gene-analysis set comprised 40 patients (16 LN1+ and 24 LN0), with one observation per patient after tissue-core curation and quality control. The 42-patient availability count is used only for cohort and TMA-image accounting; internal differential-expression, score, AUROC, and association analyses used the curated 40-patient set.

H&E slides were digitized at a nominal source resolution of 0.25 micrometers per pixel (40x equivalent). The implemented 20x workflow extracted non-overlapping 624 x 624-pixel source tiles, resized them to 312 x 312 pixels for tissue-content and color-normalization processing, and subsequently resized retained tiles to 224 x 224 pixels for encoder input. Candidate tissue cores were reviewed under the documented pathology and image-quality procedure.

### Tile eligibility

A Sobel-gradient magnitude image was computed for each candidate tile. Let *G_p_* be the gradient magnitude at pixel *p*, *τ_G_* = 15 the edge threshold, and *P* the tile-pixel set. A tile was retained when at least half of its pixels exceeded the threshold:

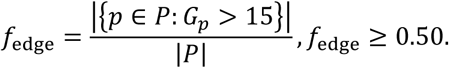

Retained tiles underwent Macenko stain normalization^48^ and were subsequently resized to 224 × 224 pixels for the encoder.

### Path2Space tile-level expression inference

Each normalized tile was standardized using the ImageNet channel means and standard deviations, then encoded with CTransPath^31^ to a 768-dimensional feature vector. The inference model applied a multilayer perceptron (MLP) to each tile independently:

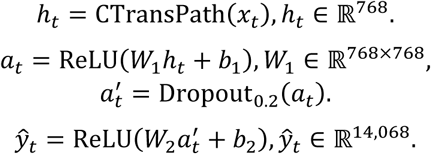

The tile-level vector *y_t_* contains image-derived estimates for 14,068 genes. For a core *c* with eligible tile set *T_c_*, core-level expression used an unweighted arithmetic mean:

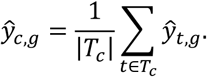

The image-derived estimates were used to nominate candidates and characterize internal associations. They were not treated as direct measurements of RNA abundance. Because the gene-specific image-to-expression correlations for TMEM184A and ZNF57 were weak, the defensible interpretation is: Path2Space-derived image-based expression estimates nominated TMEM184A and ZNF57, whose associations were subsequently evaluated using measured transcriptomic data in independent cohorts.

### Internal patient-level differential-expression analysis

Differential-expression testing compared the curated 16 LN1+ and 24 LN0 patients across the 14,068 Path2Space output genes^30^. For gene *g*, the log fold change used a small stabilizing constant *ε* = 10^−9^:

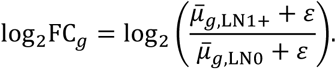

A two-sided Welch t test allowed unequal variances. If *x̅*_1*g*_, *s*_1*g*_^2^, and *n*_1_ denote the LN1+ mean, variance, and sample size and the subscript 0 denotes LN0, then:

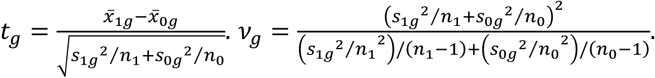

Benjamini-Hochberg (BH)^49^ adjustment controlled the false-discovery rate across finite tests. Of 14,068 genes, 13,870 yielded finite tests and 198 were undefined; 118 had nominal P < 0.05, but none had q < 0.05 (minimum q = 0.93). Thus the 118-gene set is explicitly a nominal, TMA-anchored candidate set rather than a false-discovery-rate-significant signature.

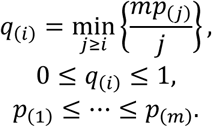

Descriptive uncertainty intervals shown in supplementary analyses used 4,000 stratified patient bootstrap resamples. These intervals do not change the nominal feature-selection rule.

### External transcriptomic cohorts and phenotype harmonization

Independent measured-transcriptomic cohorts were used to evaluate direction and nodal association without refitting graph-module membership to the external outcome. FUSCC expression was analyzed as log2(FPKM + 1). The SCAN-B initiative has been described previously^50^. The supplied TCGA and SCAN-B matrices were already supplied on a log-scale and were not re-transformed by the cohort loader. Cohort-specific expression processing, nodal-endpoint definitions, and verified sample counts are summarized in **Table 2**.

**Table 2.**
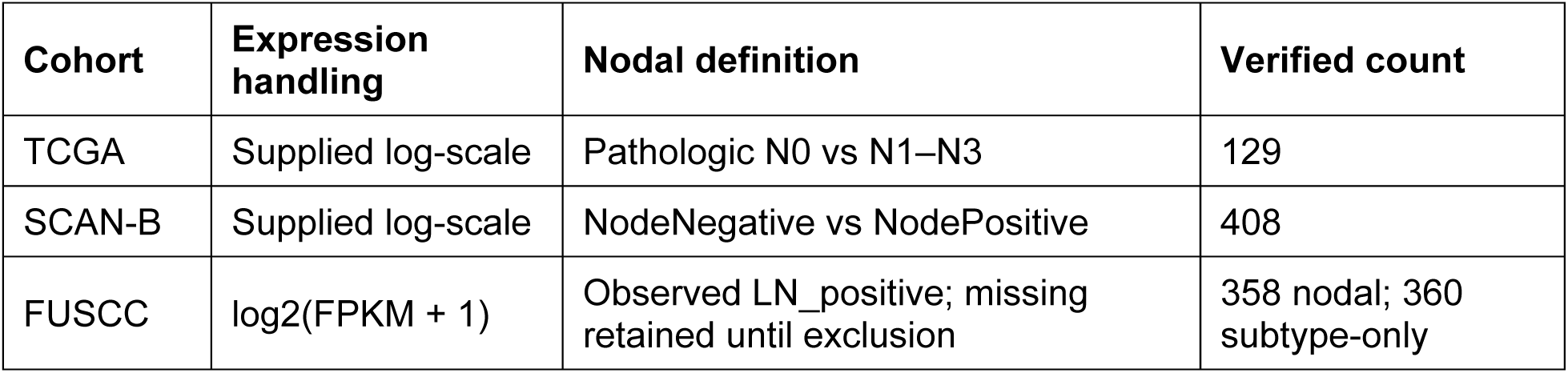
Expression handling and nodal-endpoint harmonization.

Two FUSCC records without observed nodal outcome were excluded from nodal models. They were not coerced to LN0. METABRIC and the France cohort were not used in the primary nodal validation because treatment-naïve status was not confirmed and, for the France cohort, the available endpoint was distant metastasis rather than regional nodal involvement.

### Cross-cohort candidate-gene nomination

The internal nominal candidate set comprised the 118 genes with Welch-test P < 0.05. For each candidate, the direction of the LN1+ versus LN0 effect and nominal cohort-specific significance were evaluated in the internal TMA and three independent measured-transcriptomic cohorts. A gene was considered directionally concordant across all four cohorts when the sign of its cohort-specific effect agreed in every cohort. Thirteen genes met the all-cohort direction criterion; two genes, TMEM184A and ZNF57, were also nominally significant in all four.

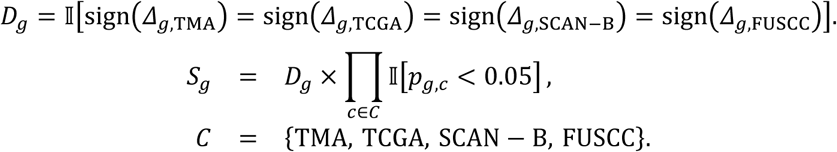

### TNBC graph construction from bulk and single-cell data

A TNBC-specific co-expression graph was constructed exclusively from measured transcriptomic data, in which genes were represented as nodes and retained gene-gene co-expression relationships as edges; the active graph did not incorporate STRING or other prior interation databases. The implemented sequence was: cohort-specific bulk candidate-edge nomination, sign-consistent bulk consensus, epithelial-cell restriction and single-cell candidate-edge nomination, weighted evidence fusion, largest-connected-component restriction, spectral embedding, row normalization, and cosine-neighborhood extraction. Bulk evidence came from TCGA (n = 129), SCAN-B (n = 419 for graph construction), and FUSCC (n = 360 for graph construction). Single-cell evidence came from epithelial cells in the Karaayvaz^32^, Wu^33^, and Pal^34^ atlases. Graph-construction counts differ from nodal-model counts because expression availability, rather than a non-missing nodal endpoint, determined graph inclusion.

### Bulk-cohort candidate edges

Within each bulk cohort c, Spearman rank correlation ρ_ijc was computed between every eligible pair of genes i and j. For each focal gene i, only its 30 partners with largest absolute correlation were retained, subject to |ρ_ijc |≥0.30. An undirected candidate pair was considered supported by cohort c if it appeared in either directed top-30 list:

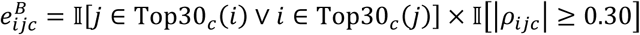

Bulk consensus additionally required the correlation sign to agree in at least two of three cohorts. Let *B_ij_* be the integer number of supporting bulk cohorts after this sign-consistency rule; the normalized bulk support was 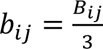. The resulting bulk consensus contained 154,177 edges:

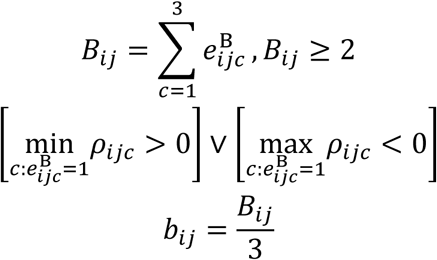

### Single-cell epithelial restriction and preprocessing

Cells were classified as epithelial when EPCAM was detected or at least two of KRT8, KRT18, and KRT19 were detected. For cell *k* with raw count *x_gk_* for gene *g* and library size *L_k_*, expression was counts-per-10,000 normalized and log transformed:

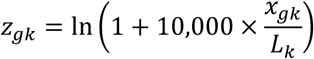

Genes had to be detected in at least 30 epithelial cells. To control computational load, at most 15,000 epithelial cells were included per atlas. Spearman correlations were computed through average ranks to handle tied expression values. As in bulk data, each gene retained its 30 largest-absolute correlation partners, now requiring 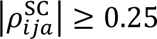 for atlas *a*. The three atlas-specific edge counts were 593^32^, 12,089^33^, and 15,000^34^ (capped).

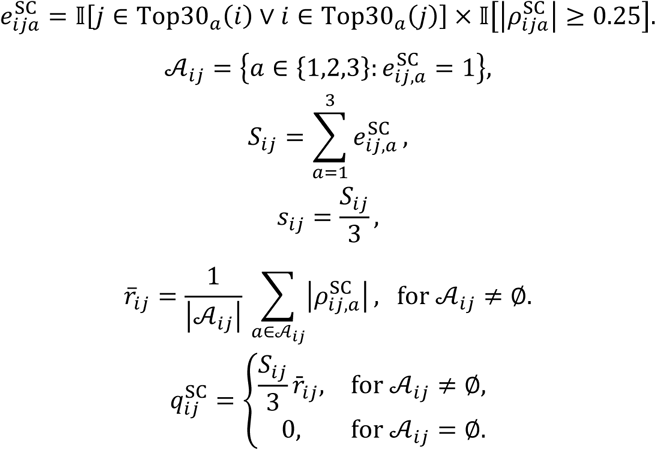

### Evidence fusion

Bulk and single-cell evidence were merged into a non-negative edge confidence *c_ij_*. The active implementation gives unit weight to normalized bulk support and a 0.6 multiplier to normalized single-cell support, modulated by the mean absolute single-cell correlation:

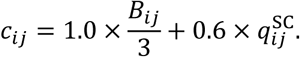

Pairs supported only by one evidence source could enter through the corresponding non-zero term. Duplicate undirected pairs were collapsed to a single weighted edge. The merged graph contained 513,122 edges.

### Spectral graph embedding and module definition

#### Weighted adjacency and connected component

Let *V* be the graph’s gene set and *A* its symmetric weighted adjacency matrix, with *A_ij_* = *c_ij_* for a retained edge and *A_ij_* = 0 otherwise. Self-edges were not used. The degree matrix *D* is diagonal:

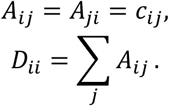

Spectral embedding was restricted to the largest connected component, which contained 12,609 genes from the 14,068-gene Path2Space universe.

### Symmetric normalized adjacency and eigendecomposition

The implementation eigendecomposed the symmetric normalized adjacency *P*, which is equivalent to working with the symmetric normalized Laplacian *L*_sym_ = *I* - *P*:

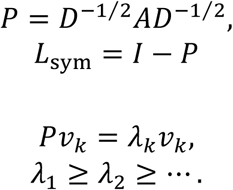

The leading trivial eigenvector was removed. The next *d* = 200 eigenvectors formed a matrix *E* with |*V*| rows and 200 columns. Each gene row was normalized to unit Euclidean length:

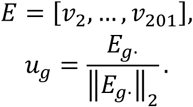

### Cosine neighborhoods

Graph proximity between genes *g* and ℎ was defined as the cosine similarity of their row-normalized spectral coordinates. Because rows were normalized, cosine similarity is their inner product:

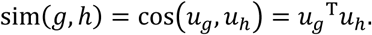

For a hub gene ℎ, its graph neighborhood excluded the hub itself and retained genes with similarity at least 0.50:

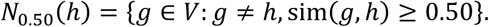

The active neighborhood tables contain 100 TMEM184A neighbors (minimum retained cosine 0.5148302) and 52 ZNF57 neighbors (minimum retained cosine 0.50010264). The score constructors add the hub to its neighbor list, producing 101-gene and 53-gene scores, respectively. The hub is therefore included in AUC, odds-ratio, and active module-score analyses. In contrast, the enrichment analysis used only the 100 or 52 neighbors to Enrichr and does not include the hub.

Two- and three-dimensional t-SNE or UMAP coordinates are visualization layers only. They do not define active module membership, which is determined in the 200-dimensional spectral space by the cosine threshold above.

### Graph robustness and biological-recovery analyses

#### Embedding-dimension selection

Embedding dimension was evaluated by reconstructing held-out graph edges from pairwise spectral similarity. Reported edge-reconstruction AUROCs were 0.8527 at 100 dimensions, 0.8549 at 200 dimensions, and 0.8595 at 350 dimensions. The 200-dimensional embedding was selected as a plateau compromise rather than as the numerical maximum.

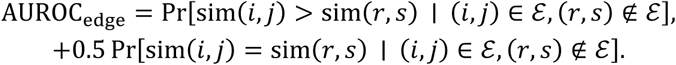

#### Edge-bootstrap neighborhood stability

The merged edge list was resampled with replacement 50 times. Each bootstrap graph was independently embedded, and a fixed-k = 100 neighborhood was recovered for each target hub. Pairwise Jaccard similarity was evaluated across the 50 bootstrap replicates, giving 1,225 pairwise replicate comparisons. These comparisons share bootstrap replicates and therefore summarize stability rather than independent inferential replicates:

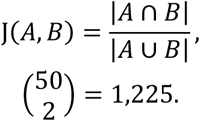

#### Similarity-cutoff sensitivity

The empirical random-pair inclusion rate at cosine 0.50 was 0.61%. Cutoffs from 0.30 through 0.70 were swept. Across this range, the TMEM184A module’s nodal AUROC remained approximately 0.595–0.603, and leave-one-cohort-out values remained approximately 0.587–0.601. Cutoff sensitivity is a robustness analysis, not an outcome-driven cutoff optimization.

#### Bulk-only recovery

The full multi-source graph neighborhoods were compared with neighborhoods reconstructed from bulk evidence only. Jaccard recovery was 0.60 for TMEM184A and 0.74 for ZNF57; analytic chance expectations were 0.0036 and 0.0018, respectively.

#### Recovery of biological structure

Four pre-specified biological gene sets were evaluated against 2,000 degree-matched random sets. Empirical one-sided P values used a plus-one correction:

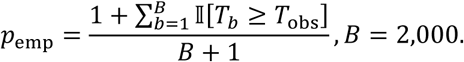

Each of the four sets yielded pemp = 0.0005, the minimum possible under this correction. In a pathway-level recovery analysis, 1,879 Reactome pathways of size 5–200 were evaluated, with at most 60 within-pathway pairs per pathway and 20,000 positive pairs matched to 20,000 random pairs. The observed AUROC was 0.533. Coordinate-label permutation (200 permutations) yielded an empirical P = 0.005. A filtered analysis removed direct known interactions.

### Gene-score construction and performed scaling

#### Cohort-internal gene standardization

Within each analyzed cohort, expression was standardized separately for each gene using the cohort arithmetic mean and the standard deviation convention implemented by the corresponding analysis script. For sample i, gene g, and cohort c:

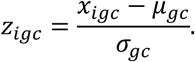

#### Two-gene score

We developed the following formula for computing the two-gene score for sample i in cohort c:

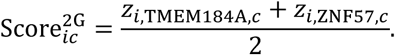

z_i,g,c is within-cohort standardized expression; TMEM184A and ZNF57 are gene labels; Score^2G_ic^ is their arithmetic mean.

#### Hub-inclusive module scores

Let *M*_ℎ_ contain the hub ℎ together with all genes in *N*_0.50_(ℎ). The active module score is the unweighted mean of within-cohort standardized expression over the hub-inclusive constructor:

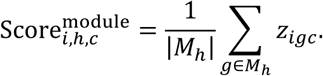

The gene sets used to construct the two-gene score, hub-inclusive module scores, and pathway-enrichment inputs are summarized in **Table 3**.

**Table 3.**
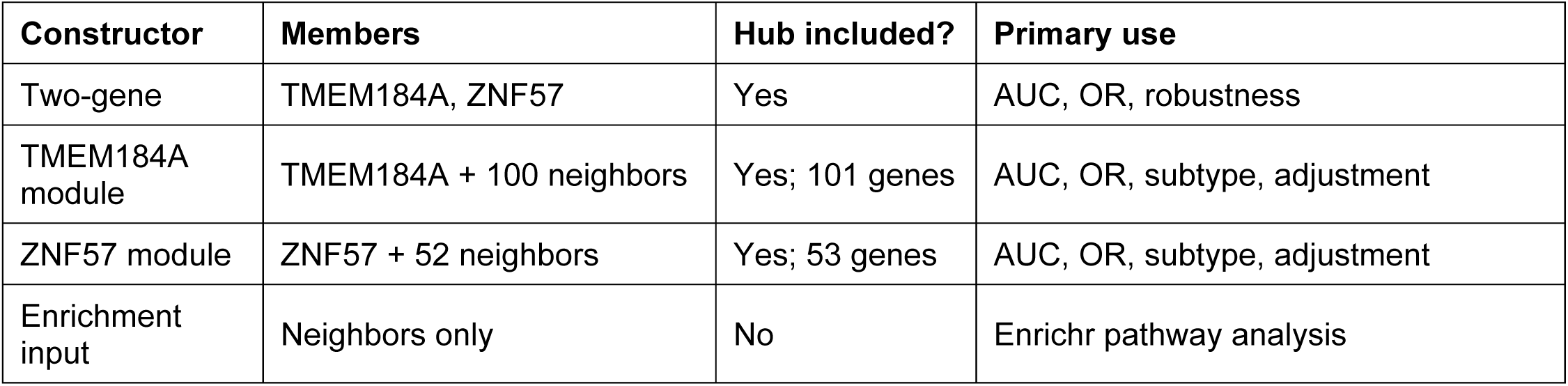
Active gene-score constructors.

| **Constructor** | **Members** | **Hub included?** | **Primary use** |
|---|---|---|---|
| Two-gene | TMEM184A, ZNF57 | Yes | AUC, OR, robustness |
| TMEM184A module | TMEM184A + 100 neighbors | Yes; 101 genes | AUC, OR, subtype, adjustment |
| ZNF57 module | ZNF57 + 52 neighbors | Yes; 53 genes | AUC, OR, subtype, adjustment |
| Enrichment input | Neighbors only | No | Enrichr pathway analysis |

For odds-ratio reporting, each composite score was standardized again within the analysis cohort so that the coefficient represented a one-standard-deviation increase. Module weights and membership were not refit to nodal outcomes in external cohorts.

### Nodal-association models and meta-analysis

#### Discrimination

AUROC quantified separation of LN1+ from LN0. For a continuous score *S*, AUROC has the probability interpretation:

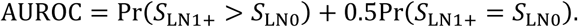

Patient-level bootstrap intervals were used where reported. Ties were handled by the standard rank-based AUROC convention. No clinical operating threshold was optimized for primary external validation.

#### Cohort-specific logistic regression

For cohort *c*, unpenalized logistic regression related observed nodal status *Y_ic_* to a standardized score

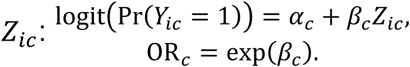

Standard errors were taken from the inverse observed Fisher information; Wald statistics produced two-sided P values and 95% confidence intervals

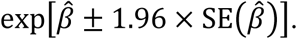

#### Random-effects pooling

Cohort log odds ratios *y_c_* = log(OR*_c_*) with within-study variances *v_c_* were combined using the DerSimonian–Laird estimator^51^. With fixed-effect weights 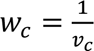, Cochran’s *Q* and between-study variance *τ*^2^ were:

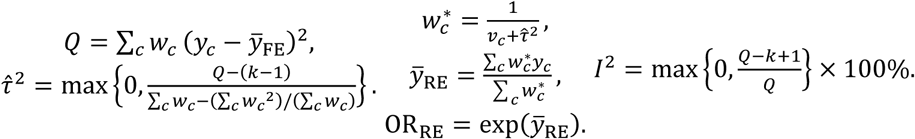

#### Score quartiles

Patients were assigned to cohort-specific score quartiles using empirical quantile binning. The LN1+ proportion in each quartile received a Wilson 95% confidence interval. A Cochran–Armitage trend test evaluated ordered change across quartiles. Quantile analyses are descriptive complements to the continuous-score model.

#### Leave-one-cohort-out discrimination

For external leave-one-cohort-out evaluation, model fitting excluded the external cohort being tested, frozen model coefficients were applied to that held-out cohort, and AUROC was calculated from held-out patient scores. The internal TMA discovery cohort was displayed separately. Patient-bootstrap 95% confidence intervals were computed in the held-out cohort; bootstrap samples containing only one outcome class were not used.

### TNBC subtype analyses

Published TNBCtype-4 calls were used without local relabeling where available. TCGA included 118 samples: BL1 n = 42, BL2 n = 22, M n = 28, and LAR n = 26, based on the Lehmann 2016 subtype table^5^. FUSCC subtype-only analysis included 360 samples: BLIS n = 139, IM n = 87, LAR n = 81, and MES n = 53.

For each score or gene feature, overall subtype differences were tested by the Kruskal–Wallis rank test. Each subtype was also compared one-versus-rest by AUROC and the Mann–Whitney U test. Cliff’s delta was derived from AUROC:

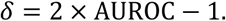

Patient-level bootstrap confidence intervals used 2,000 resamples. BH correction was applied as one family across 40 one-versus-rest comparisons (2 cohorts x 5 features x 4 subtype contrasts). Supplementary luminal/androgen-receptor composite correlations were analyzed using Spearman correlation, Fisher-transformation 95% confidence intervals, and BH correction across 15 cohort-by-feature correlation tests.

#### Within-subtype nodal discrimination

Within each available subtype, nodal discrimination of the TMEM184A module was summarized by AUROC with patient-bootstrap 95% confidence intervals. Up to 5,000 bootstrap resamples were drawn per subgroup; resamples containing only one nodal-outcome class were discarded. Two-sided Mann-Whitney U tests were adjusted by BH across the eight cohort-by-subtype tests. FUSCC used native Fudan/Jiang subtype calls and TCGA used published Lehmann TNBCtype-4 calls; similarly named categories were not treated as interchangeable across systems.

### Functional enrichment and module characterization

#### Over-representation analysis

The Enrichr/gseapy workflow^46,47^ submitted graph neighbors only—100 for TMEM184A and 52 for ZNF57—rather than the hub-inclusive score constructors. Five libraries were queried: MSigDB Hallmark 2020^52^, Reactome Pathways 2024^53^, KEGG^54^ 2021 Human, WikiPathways^55^ 2024 Human, and GO Biological Process^56^ 2026. Enrichr-adjusted P values were reported using BH correction.

#### Cross-cell module coupling

In the single-cell independence analysis, per-cell module scores were computed in each of three independent TNBC single-cell atlases after restriction to cancer epithelial cells. Pearson correlation between the two module scores, *r*, was compared with the retained 95% interval from size-matched random gene sets evaluated in the same cells. Let *T* and *Z* be the two per-cell module-score vectors:

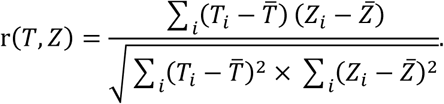

#### Within-module coherence and noise robustness

Within-module coherence was evaluated separately in TCGA, SCAN-B, and FUSCC as the mean absolute Pearson correlation across all distinct gene pairs in each hub-inclusive module. Each observed value was compared with size-matched random gene sets sampled without replacement from genes shared across all three cohorts, and empirical upper-tail P values used a plus-one correction. For noise robustness, zero-mean Gaussian noise was added to the FUSCC gene-standardized expression matrix at 11 standard-deviation levels from 0 to 2 in increments of 0.2. At each level, 40 perturbations were generated using random-state seed 0, and the mean Spearman correlation between the perturbed and unperturbed patient rankings was calculated for the module score and hub gene. Split-half module reliability was averaged across 50 random gene splits and corrected with the Spearman-Brown formula.

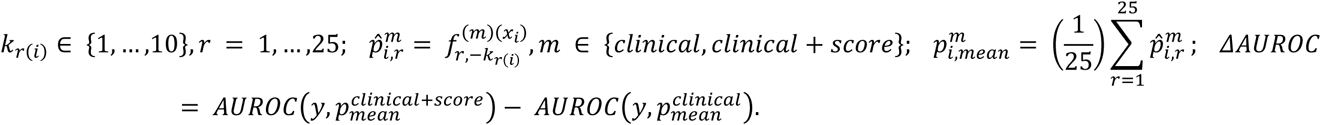

### Clinical-variable comparison and adjusted sensitivity models

#### Paired cross-validated clinical comparison

The Supplementary Figure S6D analysis used a paired cross-validated AUROC comparison. Clinical-only and clinical-plus-score logistic models used 10-fold stratified cross-validation repeated across 25 seeds. Out-of-fold probabilities were averaged per patient across repeats, and the AUROC difference was evaluated with 5,000 paired patient bootstrap resamples. Complete-case counts were FUSCC n = 323, SCAN-B n = 394, and TCGA n = 75. Clinical predictors were age, tumor size, and grade for FUSCC and SCAN-B, and age plus pathologic T category for TCGA.

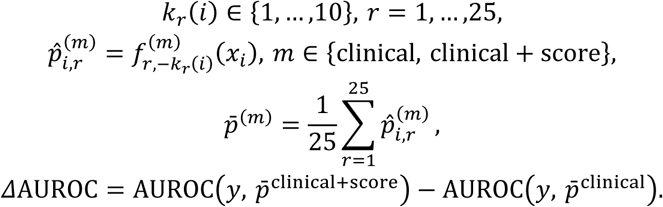

#### Expression-derived proliferation and purity sensitivity covariates

Adjustment analyses treated proliferation and the expression-derived purity proxy as sensitivity covariates. The detailed FUSCC complete-case analysis used n = 287 patients, including 112 LN1+ patients; 71 of the 358 records with observed nodal outcome were excluded by complete-case requirements. The purity proxy was not interpreted as a pathological purity assessment or as an ABSOLUTE- or ESTIMATE-derived purity value.

The proliferation covariate was the unweighted mean of within-cohort standardized expression for 19 cell-cycle genes: MKI67, CCNB1, CCNB2, CDK1, TOP2A, BIRC5, AURKA, PLK1, CENPF, UBE2C, RRM2, TYMS, CCNA2, CDC20, KIF11, NUSAP1, PCNA, FOXM1, and TPX2.

The purity proxy was constructed from the same cohort expression matrix. The initial epithelial marker set was EPCAM, KRT8, KRT18, CDH1, KRT19, and KRT7; KRT19 and KRT7 were removed because they belonged to an active tested module, leaving EPCAM, KRT8, KRT18, and CDH1. Stromal markers were COL1A1, COL1A2, COL3A1, COL5A1, FAP, THY1, PDGFRB, SPARC, FN1, and VIM. Immune markers were PTPRC, CD3D, CD2, CD8A, LCK, CD53, CD37, CCL5, GZMA. Each marker was standardized within cohort before the epithelial, stromal, and immune means were calculated. No retained purity marker overlapped an active module constructor.

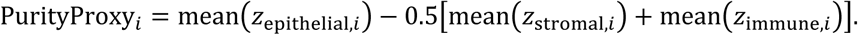

The proxy was used only to assess sensitivity to expression patterns compatible with specimen composition. Because the proxy and module scores were derived from the same expression matrix, adjusted estimates were interpreted as sensitivity analyses rather than causal or clinical models.

#### Adjusted logistic-regression specifications

For the current single-covariate adjustment analysis, each standardized score entered an unpenalized logistic model alone and then with either the proliferation score or the expression-derived purity proxy. Odds ratios were reported per one-standard-deviation increase in the score. The displayed collection contained 54 exploratory estimates; nominal two-sided Wald P values were not adjusted across this sensitivity family.

The detailed matched-sample FUSCC sensitivity series used a common complete-case set across nested specifications: score only; score plus molecular-subtype indicators; score plus proliferation; score plus purity proxy; score plus clinicopathologic covariates; a full model containing subtype, proliferation, purity, histological grade, log-transformed positive tumor size, age, and Ki-67; and the full model refitted after excluding LAR tumors. Continuous covariates were standardized before fitting; binary or categorical indicator columns were not standardized.

Models were fit with scikit-learn LogisticRegression using no penalty, the lbfgs solver, and a maximum of 8,000 iterations. Standard errors were calculated from the inverse observed Fisher-information matrix. Two-sided Wald statistics produced nominal P values and 95% confidence intervals, and exponentiated coefficients were reported as odds ratios.

### Quantification and Statistical Analysis

Unless specified otherwise, analyses were patient-level and statistical tests were two-sided. Missing nodal labels were excluded from nodal analyses and were never converted to LN0. Complete-case analysis was used where covariates were required; no imputation procedure was applied in the reported adjusted models. Confidence intervals were 95%. Bootstrap resampling was performed at the patient level, and a prespecified random seed or seed sequence was used where the retained script defined one. BH correction was applied only to the explicitly defined hypothesis families described below; nominal sensitivity analyses are labeled as such. Analysis-specific statistical procedures, analysis sets, multiplicity control, and uncertainty-estimation methods are summarized in **Table 4**.

**Table 4.**
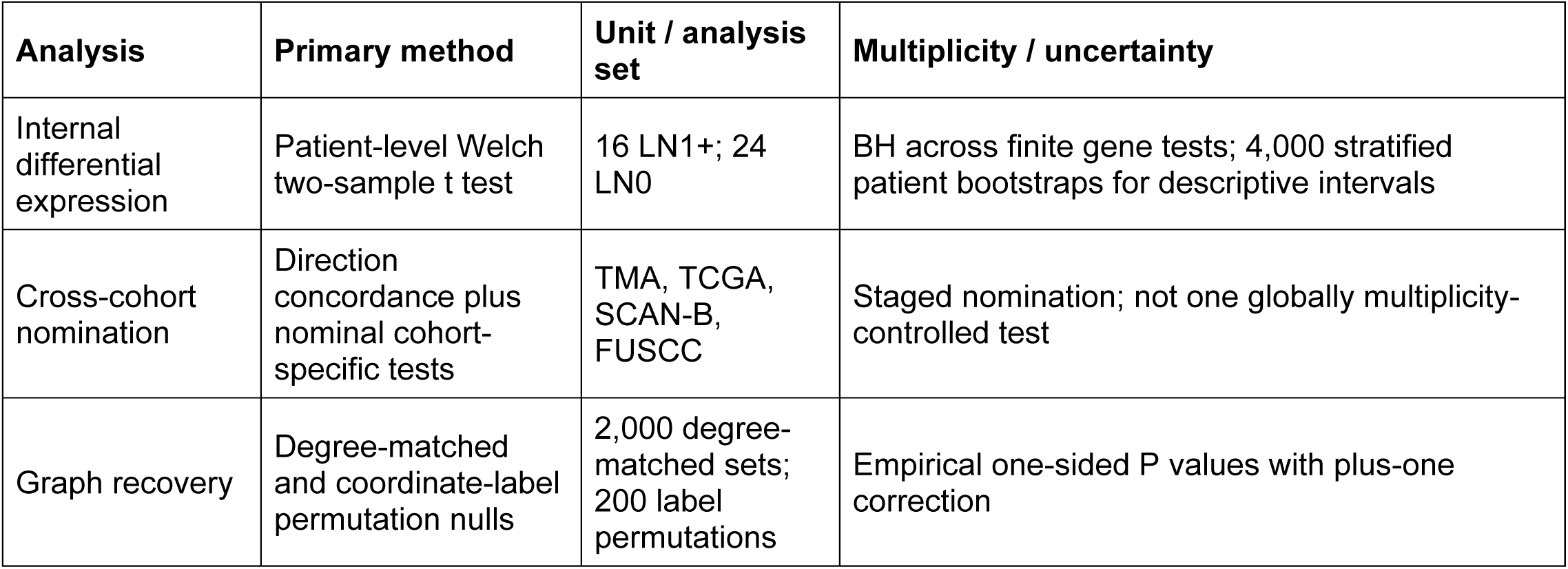
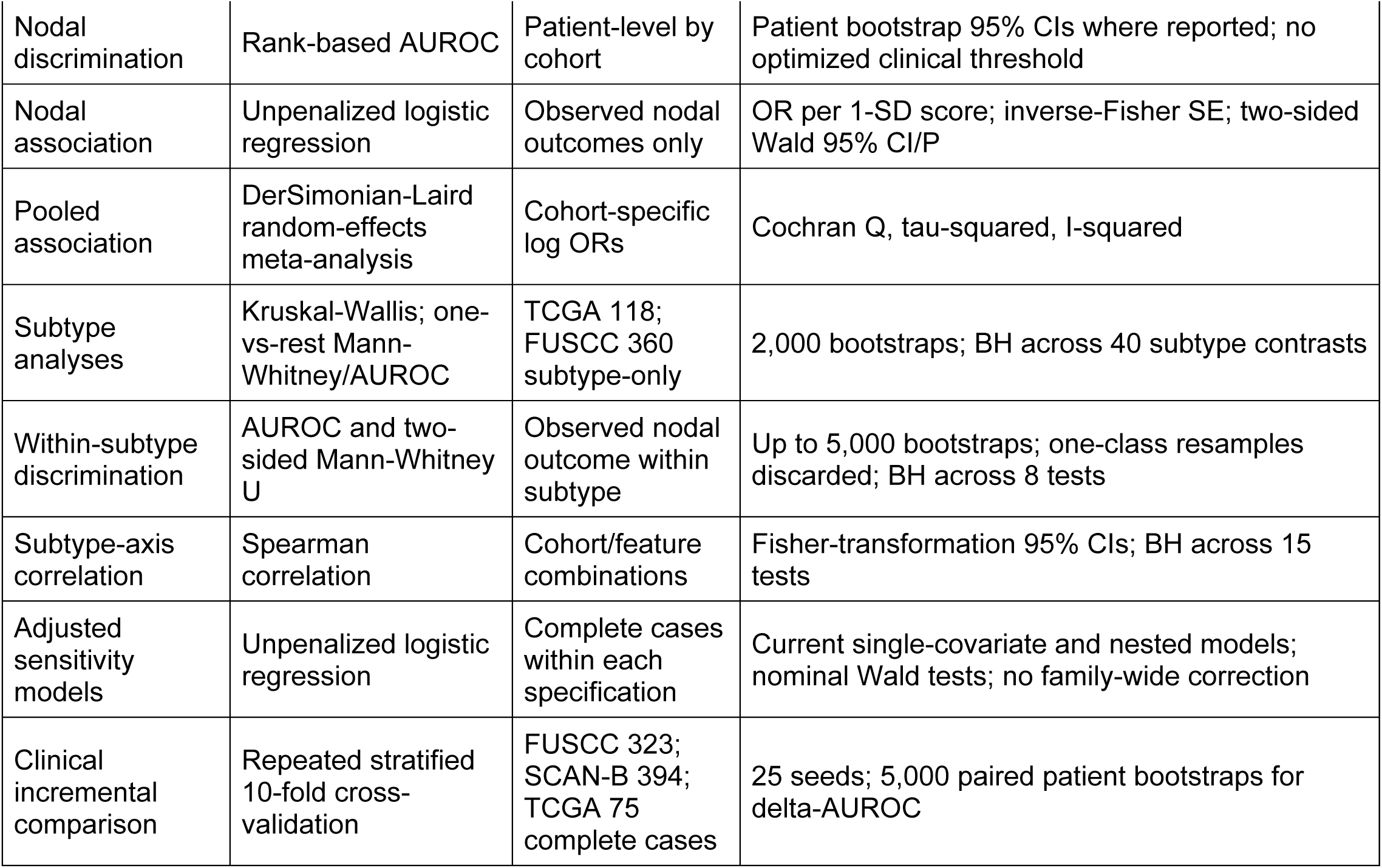
Analysis-specific statistical procedures and uncertainty reporting.

### Analysis-specific procedures

[tbl4]

### Software and reproducibility

Core numerical and tabular analyses were implemented in Python using NumPy^57^, pandas, SciPy^58^, and scikit-learn^59^. Image-expression inference used PyTorch^60^-compatible CTransPath^31^ model code. Enrichment queries used gseapy^46^ and Enrichr^47^, and manuscript figures were generated with Matplotlib^61^-based scripts.

## REFERENCES

1 Bianchini, G., Balko, J. M., Mayer, I. A., Sanders, M. E. & Gianni, L. Triple-negative breast cancer: challenges and opportunities of a heterogeneous disease. Nat Rev Clin Oncol 13, 674–690 (2016). 10.1038/nrclinonc.2016.66

2 Jiang, Y. Z. et al. Genomic and Transcriptomic Landscape of Triple-Negative Breast Cancers: Subtypes and Treatment Strategies. Cancer Cell 35, 428–440 e425 (2019). 10.1016/j.ccell.2019.02.001

3 Foulkes, W. D., Smith, I. E. & Reis-Filho, J. S. Triple-negative breast cancer. New Engl J Med 363, 1938–1948 (2010).

4 Lehmann, B. D. et al. Identification of human triple-negative breast cancer subtypes and preclinical models for selection of targeted therapies. J Clin Invest 121, 2750–2767 (2011). 10.1172/JCI45014

5 Lehmann, B. D. et al. Refinement of Triple-Negative Breast Cancer Molecular Subtypes: Implications for Neoadjuvant Chemotherapy Selection. Plos One 11 (2016). ARTN e0157368 10.1371/journal.pone.0157368

6 Denkert, C., Liedtke, C., Tutt, A. & von Minckwitz, G. Molecular alterations in triple-negative breast cancer-the road to new treatment strategies. Lancet 389, 2430–2442 (2017). 10.1016/S0140-6736(16)32454-0

7 Dent, R. et al. Triple-negative breast cancer: clinical features and patterns of recurrence. Clin Cancer Res 13, 4429–4434 (2007). 10.1158/1078-0432.CCR-06-3045

8 Anders, C. K., Zagar, T. M. & Carey, L. A. The management of early-stage and metastatic triple-negative breast cancer: a review. Hematol Oncol Clin North Am 27, 737–749, viii (2013). 10.1016/j.hoc.2013.05.003

9 Zeichner, S. B., Terawaki, H. & Gogineni, K. A Review of Systemic Treatment in Metastatic Triple-Negative Breast Cancer. Breast Cancer (Auckl) 10, 25–36 (2016). 10.4137/BCBCR.S32783

10 Karagiannis, G. S. et al. Neoadjuvant chemotherapy induces breast cancer metastasis through a TMEM-mediated mechanism. Science translational medicine 9, eaan0026 (2017).

11 Karagiannis, G. S., Condeelis, J. S. & Oktay, M. H. Chemotherapy-induced metastasis: mechanisms and translational opportunities. Clin Exp Metastasis 35, 269–284 (2018). 10.1007/s10585-017-9870-x

12 Wu, Y. J. et al. Cyclophosphamide enhances human tumor growth in nude rat xenografted tumor models. Neoplasia 11, 187–195 (2009). 10.1593/neo.81352

13 Ogier du Terrail, J., et al. Federated learning for predicting histological response to neoadjuvant chemotherapy in triple-negative breast cancer. Nat Med 29, 135–146 (2023). 10.1038/s41591-022-02155-w

14 Coudray, N. et al. Classification and mutation prediction from non-small cell lung cancer histopathology images using deep learning. Nat Med 24, 1559–1567 (2018). 10.1038/s41591-018-0177-5

15 Kather, J. N. et al. Deep learning can predict microsatellite instability directly from histology in gastrointestinal cancer. Nature medicine 25, 1054–1056 (2019).

16 Amgad, M. et al. A population-level digital histologic biomarker for enhanced prognosis of invasive breast cancer. Nature medicine 30, 85–97 (2024).

17 Vorontsov, E. et al. A foundation model for clinical-grade computational pathology and rare cancers detection. Nat Med 30, 2924–2935 (2024). 10.1038/s41591-024-03141-0

18 Ståhl, P. L. et al. Visualization and analysis of gene expression in tissue sections by spatial transcriptomics. Science 353, 78–82 (2016). 10.1126/science.aaf2403

19 Moncada, R. & d Barkley, F. W. M. chiodin, J. c. devlin, M. Baron, ch hajdu, d. M. Simeone, i. Yanai, integrating microarray-based spatial transcriptomics and single-cell RnA-seq reveals tissue architecture in pancreatic ductal adenocarcinomas. Nat. Biotechnol 38, 333–342 (2020).

20 Jackson, H. W. et al. The single-cell pathology landscape of breast cancer. Nature 578, 615–620 (2020). 10.1038/s41586-019-1876-x

21 Danenberg, E. et al. Breast tumor microenvironment structures are associated with genomic features and clinical outcome. Nat Genet 54, 660–669 (2022). 10.1038/s41588-022-01041-y

22 Wang, X. Q. et al. Spatial predictors of immunotherapy response in triple-negative breast cancer. Nature 621, 868–876 (2023). 10.1038/s41586-023-06498-3

23 Shiao, S. L. et al. Single-cell and spatial profiling identify three response trajectories to pembrolizumab and radiation therapy in triple negative breast cancer. Cancer Cell 42, 70–84 e78 (2024). 10.1016/j.ccell.2023.12.012

24 Klughammer, J. et al. A multi-modal single-cell and spatial expression map of metastatic breast cancer biopsies across clinicopathological features. Nat Med 30, 3236–3249 (2024). 10.1038/s41591-024-03215-z

25 Caire, R. et al. A 3D morphogenetic blueprint for metastatic outgrowth in breast cancer. Cell 189, 3701–3718 e3730 (2026). 10.1016/j.cell.2026.03.009

26 Gan, S. et al. Distinct tumor architectures and microenvironments for the initiation of breast cancer metastasis in the brain. Cancer Cell 42, 1693–1712 e1624 (2024). 10.1016/j.ccell.2024.08.015

27 Chen, W. et al. A visual-omics foundation model to bridge histopathology with spatial transcriptomics. Nat Methods 22, 1568–1582 (2025). 10.1038/s41592-025-02707-1

28 Huang, T., Liu, T., Babadi, M., Ying, R. & Jin, W. STPath: a generative foundation model for integrating spatial transcriptomics and whole-slide images. NPJ Digit Med 8, 659 (2025). 10.1038/s41746-025-02020-3

29 Wang, C. et al. Benchmarking the translational potential of spatial gene expression prediction from histology. Nat Commun 16, 1544 (2025). 10.1038/s41467-025-56618-y

30 Shulman, E. D. et al. AI-predicted spatial transcriptomics unlocks breast cancer biomarkers from pathology. Cell (2026).

31 Wang, X. et al. Transformer-based unsupervised contrastive learning for histopathological image classification. Med Image Anal 81, 102559 (2022). 10.1016/j.media.2022.102559

32 Karaayvaz, M. et al. Unravelling subclonal heterogeneity and aggressive disease states in TNBC through single-cell RNA-seq. Nat Commun 9, 3588 (2018).

33 Wu, S. Z. et al. A single-cell and spatially resolved atlas of human breast cancers. Nat Genet 53, 1334–1347 (2021). 10.1038/s41588-021-00911-1

34 Pal, B. et al. A single-cell RNA expression atlas of normal, preneoplastic and tumorigenic states in the human breast. EMBO J 40, e107333 (2021). 10.15252/embj.2020107333

35 Song, A. H. et al. Artificial intelligence for digital and computational pathology. Nat Rev Bioeng 1, 930–949 (2023). 10.1038/s44222-023-00096-8

36 Wang, X. et al. A pathology foundation model for cancer diagnosis and prognosis prediction. Nature 634, 970–978 (2024). 10.1038/s41586-024-07894-z

37 Saltz, J. et al. Spatial Organization and Molecular Correlation of Tumor-Infiltrating Lymphocytes Using Deep Learning on Pathology Images. Cell Rep 23, 181–193 e187 (2018). 10.1016/j.celrep.2018.03.086

38 Yang, J. et al. Guidelines and definitions for research on epithelial-mesenchymal transition. Nat Rev Mol Cell Biol 21, 341–352 (2020). 10.1038/s41580-020-0237-9

39 Ecco, G., Imbeault, M. & Trono, D. KRAB zinc finger proteins. Development 144, 2719–2729 (2017). 10.1242/dev.132605

40 Marx, V. Method of the Year: spatially resolved transcriptomics. Nat Methods 18, 9–14 (2021). 10.1038/s41592-020-01033-y

41 Moses, L. & Pachter, L. Publisher Correction: Museum of spatial transcriptomics. Nat Methods 19, 628 (2022). 10.1038/s41592-022-01494-3

42 Chen, R. J. et al. Towards a general-purpose foundation model for computational pathology. Nat Med 30, 850–862 (2024). 10.1038/s41591-024-02857-3

43 Lu, M. Y. et al. A visual-language foundation model for computational pathology. Nat Med 30, 863–874 (2024). 10.1038/s41591-024-02856-4

44 Xiao, Y. et al. Multi-Omics Profiling Reveals Distinct Microenvironment Characterization and Suggests Immune Escape Mechanisms of Triple-Negative Breast Cancer. Clin Cancer Res 25, 5002–5014 (2019). 10.1158/1078-0432.CCR-18-3524

45 Cancer Genome Atlas, N. Comprehensive molecular portraits of human breast tumours. Nature 490, 61–70 (2012). 10.1038/nature11412

46 Fang, Z., Liu, X. & Peltz, G. GSEApy: a comprehensive package for performing gene set enrichment analysis in Python. Bioinformatics 39 (2023). 10.1093/bioinformatics/btac757

47 Kuleshov, M. V. et al. Enrichr: a comprehensive gene set enrichment analysis web server 2016 update. Nucleic Acids Res 44, W90–97 (2016). 10.1093/nar/gkw377

48 Macenko, M. et al. A Method for Normalizing Histology Slides for Quantitative Analysis. I S Biomed Imaging, 1107-+ (2009). 10.1109/Isbi.2009.5193250

49 Benjamini, Y. & Hochberg, Y. Controlling the False Discovery Rate - a Practical and Powerful Approach to Multiple Testing. J Roy Stat Soc B 57, 289–300 (1995). 10.1111/j.2517-6161.1995.tb02031.x

50 Saal, L. H. et al. The Sweden Cancerome Analysis Network - Breast (SCAN-B) Initiative: a large-scale multicenter infrastructure towards implementation of breast cancer genomic analyses in the clinical routine. Genome Med 7, 20 (2015). 10.1186/s13073-015-0131-9

51 DerSimonian, R. & Laird, N. Meta-analysis in clinical trials. Control Clin Trials 7, 177–188 (1986). 10.1016/0197-2456(86)90046-2

52 Liberzon, A. et al. The Molecular Signatures Database (MSigDB) hallmark gene set collection. Cell Syst 1, 417–425 (2015). 10.1016/j.cels.2015.12.004

53 Milacic, M. et al. The Reactome Pathway Knowledgebase 2024. Nucleic Acids Res 52, D672–D678 (2024). 10.1093/nar/gkad1025

54 Kanehisa, M., Furumichi, M., Sato, Y., Ishiguro-Watanabe, M. & Tanabe, M. KEGG: integrating viruses and cellular organisms. Nucleic Acids Res 49, D545–D551 (2021). 10.1093/nar/gkaa970

55 Martens, M. et al. WikiPathways: connecting communities. Nucleic Acids Res 49, D613–D621 (2021). 10.1093/nar/gkaa1024

56 Gene Ontology, C., et al. The Gene Ontology knowledgebase in 2023. Genetics 224 (2023). 10.1093/genetics/iyad031

57 Harris, C. R. et al. Array programming with NumPy. Nature 585, 357–362 (2020). 10.1038/s41586-020-2649-2

58 Virtanen, P. et al. SciPy 1.0: fundamental algorithms for scientific computing in Python. Nat Methods 17, 261–272 (2020). 10.1038/s41592-019-0686-2

59 Pedregosa, F. et al. Scikit-learn: Machine learning in Python. the Journal of machine Learning research 12, 2825–2830 (2011).

60 Paszke, A. et al. Pytorch: An imperative style, high-performance deep learning library. Advances in neural information processing systems 32 (2019).

61 Hunter, J. D. Matplotlib: A 2D graphics environment. Comput Sci Eng 9, 90–95 (2007). 10.1109/Mcse.2007.55

